# Sophorolipid Candidates Demonstrate Cytotoxic Efficacy Against 2D And 3D Breast Cancer Models

**DOI:** 10.1101/2022.09.01.506226

**Authors:** Cassandra L. Roberge, Rebecca T. Miceli, Lillian R. Murphy, David M. Kingsley, Richard A. Gross, David T. Corr

## Abstract

Sophorolipids are biosurfactants derived from the non-pathogenic yeast *Starmerella bombicola*, with potential efficacy in anti-cancer applications. Simple and cost-effective synthesis of these drugs makes them a promising alternative to traditional chemotherapeutics, pending their success in preliminary drug-screening. Drug screening typically utilizes 2D cell monolayers due to their simplicity and potential for high-throughput assessment. However, 2D assays fail to capture the complexity and 3D context of the tumor microenvironment, and have consequently been implicated in the high percentage of drugs investigated *in vitro* that later fail in clinical trials. We screened two sophorolipid candidates and clinically-used chemotherapeutic, doxorubicin, on *in vitro* breast cancer models ranging from 2D monolayers to 3D spheroids, employing Optical Coherence Tomography to confirm these morphologies. We calculated corresponding IC_50_ values for these drugs and found one of the sophorolipids to have comparable toxicities to the chemotherapeutic control. Our findings show increased drug resistance associated with model dimensionality, such that all drugs tested showed that 3D spheroids exhibited higher IC_50_ values than their 2D counterparts. These findings demonstrate promising preliminary data to support the use of sophorolipids as a more affordable alternative to traditional clinical interventions and demonstrate the importance of 3D tumor models in assessing drug response.

## INTRODUCTION

Sophorolipids (SLs) are a family of glycolipid biosurfactants produced through the fermentation of non-pathogenic yeasts. Due to their good biodegradability, low environmental toxicity, and high yielding fermentation processes, they are currently the subject of extensive research by academic and industrial researchers for a wide-range of applications including ingredients in cosmetics, detergents, and textiles ^1–3^. Recent studies into potential pharmaceutical applications have reported that sophorolipids are potential anti-cancer therapeutics as they are able to disrupt the cell cycle progression and cause apoptosis and/or necrosis in a variety of cancerous cell lines, including esophageal cancer ^4^, cervical cancer ^5^, lung cancer ^6^, and breast cancer ^1,6,7^. Additionally, some sophorolipid-derived structures have relatively low toxicity to noncancerous cell lines, reducing the likelihood of harsh side effects *in vivo* ^5,8,9^, and promoting greater cytotoxic selectivity.

Sophorolipids, produced by the yeast *Starmerella bombicola*, consist of sophorose, a disaccharide of glucose units linked β-1,2-linked that may be acetylated at the 6′- and/or 6”-positions, and a hydrophobic group that, most often, is ω-1-hydroxylated oleic acid ^10^. Hydroxylated fatty acids are β-glycosidically linked to sophorose and, the fatty acid moiety may be free (acidic SLs, ASLs) or linked to the sophorose 4” forming a macrolactone (lactonic SLs, LSLs). Often, LSL diacetylated at the 6’ and 6” positions is a large fraction (>50 mol%) of the natural sophorolipids formed ^1^ (structure is shown in Figure 1A). The fact that sophorolipids are produced by fermentation processes that give high productivity and volumetric yields (~ 2 g/L/h and ~200 g/L) presents the opportunity to use these natural structures as molecular precursors for low-cost therapeutics ^10,11^. Indeed, natural and modified sophorolipids have demonstrated a wide array of biological activities including anti-cancer, anti-microbial, antiviral, and anti-inflammatory properties ^12–16^. This study focuses on diacetylated LSL (DLSL) (**Figure 1a**) and the modified sophorolipid previously reported by our group (SL-EE-D) that is synthesized by ring-opening LSL under alkaline conditions by ethanol ^3,16^ followed by lipase-catalyzed selective diacetylation at the 6’ and 6” positions (**Figure 1b**) ^2^,^16^. While promising preliminary findings were reported for natural and modified SLs^17,18^, more research is needed to advance our understanding of *in vitro* SL structure-cytotoxicity relationships on cancer and normal cell lines, as well as the corresponding mechanisms of action.

**Figure 1:**
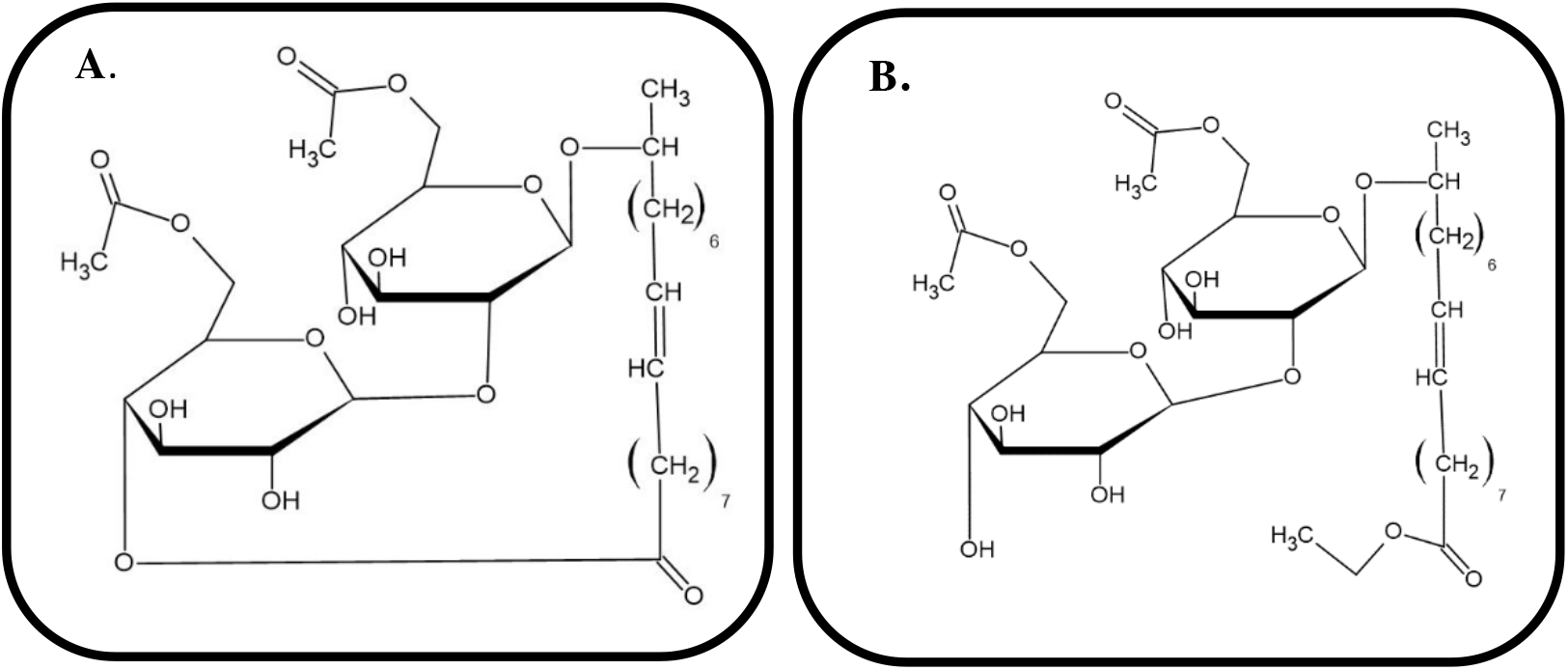
Structures of natural and modified sophorolipids studied herein: A) natural lactonic sophorolipid diacetylated at the sophorose 6’ and 6” ring positions (DLSL). B) sophorolipid ethyl ester diacetylated at the 6’ and 6” sophorose ring positions (SL-EE-D)

To test these cytotoxic activities, a suitable drug-screening platform must be employed. Traditionally, when screening for possible drug candidates in oncology, researchers turn first to *in vitro* cell models to test the proliferation inhibition and cytotoxic effects of a given drug ^19^. Investigation often begins in two-dimensional (2D) cell models, then progresses to pre-clinical *in vivo* models before moving into full clinical studies. However, prior studies have demonstrated that promising inhibitory or cytotoxicity results observed in a 2D environment are often not reflected in three-dimensional (3D) models, where increased microenvironment complexity bears a closer resemblance to physiologic conditions ^19,20^. This discrepancy has been implicated, at least in part, to the high failure rate of drug candidates during discovery, where only a low percentage of those drugs showing great promise *in vitro* succeed in clinical trials ^21^. As a result, it is now generally accepted that 3D multicellular models are more predictive of *in vivo* study outcomes than their 2D counterparts. Multicellular tumor spheroid (MCTS) cultures, in particular, exhibit several key aspects of *in vivo* tumors, such as 3D structure, metabolic function, and pathophysiological gradients, that make them an ideal screening model for new therapeutic strategies ^22,23^. Liquid overlay fabrication serves as the current gold standard for producing these models, due to its high-throughput capabilities, ease of use, and sample reproducibility ^24^. This technique utilizes non-adherent, round-bottom, 96-well plates to encourage direct cell-to-cell contact over cell-substrate contact, thus promoting cell aggregation. In cell lines with limited ability to self-aggregate, Matrigel serum (basement membrane derived from murine sarcoma) is frequently added as a supplement to liquid overlay cultures to promote formation of 3D spheroids, rather than thinner, 2.5D disk-shaped aggregates ^24^. This well documented technique is capable of producing spheroids using a variety of cell types, and serves as a popular approach for *in vitro* oncological drug-screening ^25,26^.

Triple negative human breast cancers (TNBCs) are difficult targets for drug development. In 2019, there was estimated to be over 1.7 million new cancer diagnoses and 600 thousand cancer related deaths in the US alone ^27^. TNBCs account for 15% of all breast cancers and are negative for estrogen, progesterone and HER2 hormonal receptors ^28^. This lack of hormonal receptors limits the traditional courses of treatments available. As a result, they are often approached with a combination approach of invasive surgery, radiation therapy, and especially chemotherapeutics, such as doxorubicin ^29,30^. However, these chemotherapeutics are relatively expensive to administer, which may limit their use, particularly in developing nations ^31^. Indeed, the World Health Organization noted in 2010 that in some African countries, only 20% of patients survive cancers which are curable in developed countries ^32^. Thus, the development of practical and effective treatment alternatives for TNBC patients is urgently needed.

Herein, we aim to assess the cytotoxic activity of two sophorolipid candidates on *in vitro* triple negative tumor models. Tumor models were prepared as 2D monolayers, 2.5D disk-shaped aggregates, and 3D spheroids to explore the impact of model morphology and dimensionality on drug resistance/efficacy. In addition to determining cytotoxicity on TNBC models, we determined SL toxicity against multiple non-cancerous cells to determine a therapeutic index for each compound and morphology. Additionally, the efficacy of these candidate sophorolipids was benchmarked against a gold standard chemotherapeutic – doxorubicin. By calculating the half maximal inhibitory concentration (IC_50_) for each of the tumor models, we hope to reveal possible differences in drug resistance for the three different *in vitro* tumor model morphologies, ultimately working towards both a more predictive drug-screening model and a more cost-effective alternative to traditional chemotherapeutics.

## MATERIALS AND METHODS

### Natural and modified sophorolipid (DLSL and SL-EE-D) production via fermentation

The methods for sophorolipid (SL) production have been previously described in detail ^2,3,33–35^. Briefly, LSL was synthesized through fermentation of *Starmerella bombicola* as previously reported using high oleic acid content sunflower oil as the lipid source^1^. The crude sophorolipid mixture was extracted from the fermentation broth with ethyl acetate, and the solvent was removed by rotary evaporation. Precipitation of the resulting solid in a hexane/ethyl acetate solution gave diacetylated LSL in high purity ^36^. Synthesis of SL-ethyl ester was performed as previously reported^3^, by first reacting ethanol with sodium. LSL-diacetate was treated with the sodium ethanolate solution. Upon completion, the reaction mixture was acidified, and the product was purified by recrystallization. Selective acetylation of the SL-ethyl ester primary hydroxyls at the 6’,6” positions was performed by Novozym 435 catalysis by reaction with vinyl acetate. The structures of purified lactonic sophorolipid diacetate (DLSL) and SL-ethyl ester diacetate (SL-EE-D) were confirmed by Liquid Chromatography–Mass Spectrometry (LC–MS), ^1^H-NMR, and ^13^C-NMR that are identical to those previously reported ^2,3,37^.

### Cell Culture

MDA-MB-231 triple-negative breast cancer cells (ATCC, Manassas, VA) were grown in growth medium consisting of Dulbecco’s Modified Eagle’s Medium (DMEM) supplemented with 10% (v/v) fetal bovine serum (FBS), 1% penicillin/streptomycin, and 2 mM L-glutamine. CCD-1065sk human dermal fibroblasts (HDFs) (ATCC, Manassas, VA) derived from mammary epithelial tissue were grown in growth medium consisting of DMEM supplemented with 15% (v/v) FBS and 0.5% (v/v) penicillin/streptomycin. Both cell types were cultured in standard cell culture conditions (37°C, 5% CO2, 95% RH).

### 2D Monolayer Preparation

To prepare 2D monolayer cultures ^38^, cells were trypsinized, centrifuged, and resuspended in media at 2.5 x 10^5^ cells/mL. A 100 μL volume of this cell suspension was added to each well of a 96-well flat bottom well plate, and cells were cultured for 24 hours prior to testing.

### Liquid Overlay Technique for Formation of 2.5D and 3D Tumor Aggregates

Following previously established methods ^24^, cells were trypsinized and resuspended in media at a concentration of 2.5 x 10^5^ cells/mL. 100 μL of cell suspension was added to each well of a round bottom, non-adherent, 96 well-plate (CellStar, Greiner Bio-One) to create 2.5D and 3D aggregates. To achieve 3D spheroids, these suspensions were supplemented with 2.5% Matrigel (Corning, #354263) to promote aggregation. Plates were centrifuged at 1000 rpm for 10 minutes immediately following seeding to ensure collection of the cell pellet at the bottom of each well. Plates were then incubated, and cell aggregates were cultured over a four-day maturation period.

### Aggregate morphology quantification via Optical Coherence Tomography (OCT)

Aggregate morphology was characterized using Optical Coherence Tomography (OCT) structural imaging with Imaris analysis software, as previously described ^39^. Briefly, a commercial Spectral Domain OCT (SDOCT) system, operating with a central wavelength of 1310 nm [TEL220C1, Thorlabs Inc. New Jersey, USA] was utilized, with an axial resolution of 4.2 μm (in water) and a lateral resolution of 5 μm. Image collection was performed at a 5.5kHz A-scan rate with a sensitivity of 101dB. The index of refraction of 1.33 was used for samples in liquid medium, and lateral pixel size was held constant at 1.1 μm. Aggregate morphologies were assessed using Imaris [Imaris x64, v9.5, Bitplane]. Intensity thresholding was first performed to isolate the sample region within the OCT volume scan, then aggregate outlines were traced slice-by-slice and stitched together to recreate the volume of the sample. This reconstruction was used to quantify aggregate shape characteristics, such as volume and sphericity.

### Assessment of Drug Cytotoxicity

The two candidate SLs (DLSL, SL-EE-D), as well as an established chemotherapeutic (doxorubicin) were individually solubilized in 100% DMSO before undergoing 1:2 serial dilutions in 10% DMSO. Eight concentrations were obtained for the SLs, ranging from 5000 μg/mL to 39 μg/mL. Ten concentrations of doxorubicin (Sigma, CAS 25316-40-9, lot #218824) were prepared ranging from 500 μg/mL to 7.8 μg/mL. Concentration ranges were selected to incorporate IC_50_ values seen in literature ^40–42^, and to ensure that the amount of DMSO in each well was below 1% to minimize its cytotoxic effects. Following the respective incubation periods for each model (outlined in Sections 2.3 and 2.4), 15 μL of drug and 35 μL of media were added to each well, such that the final drug concentration was 10% of the added drug concentration within each well. Three replicates were created for each concentration. Negative controls (untreated wells with cells) and positive control (media only) were included for each plate. Drugged plates were incubated for 12 hours prior to assessment.

Determination of SL drug cytotoxicity in 2D samples was performed using the cell counting kit-8 (CCK8) viability assay (Sigma Aldrich, #96992). CCK8 uses water-soluble tetrazolium to quantify the number of live cells by producing formazan dye generated by dehydrogenases in cells. Following the manufacturer’s protocol, 15 μL (one-tenth of each culture medium volume) of CCK8 was added to each well post-drug treatment, and plates were incubated for 2 hours. After incubation, 100 μL of medium from each well was transferred to a new 96-well plate. Plates were read using a BioTek uQuant Microplate Reader (BioTek Instruments) at 450 nm, with a second scan at 630 nm for background subtraction.

Assessment of SL drug cytotoxicity in 3D samples was performed using a CellTiter 3D Cell Viability Assay (Promega, #G9681) following manufacturer’s protocol. Briefly, after the 12-hour drug exposure was completed, 50 μL of cell media was removed from each well and discarded, leaving 100 μL of cells and supernatant. The plate was maintained at room temperature for 30 minutes, then 100 μL of CellTiter-Glo Luciferase detection reagent was added to each well, and the plate was placed on a shaker for 5 minutes at 500 rpm. The contents of each well were carefully pipetted into a white 96-well plate, and after 30 minutes at room temperature to allow for luminescence stabilization, the plate was imaged using a Tecan Infinite M1000 PRO Multi-Well Plate Reader, with 5 seconds of shaking prior to the scanning luminescence reading. Relative Light Units (RLU) was recorded for each well ^43^.

Doxorubicin cytotoxicity was evaluated manually for all cell models (2D, 2.5D, 3D) following previously established methods ^44^. After 12 hours of treatment, cell models were returned to cell suspensions and cells were counted via hemocytometer. To dissociate the monolayer/aggregates, 80 μL of media was removed from each well and replaced with 100 μL of trypsin. Care was taken to expose cells to trypsin for no longer than 10 minutes to reduce potential cytotoxicity associated with trypsin. After 10 minutes, wells were mixed for 30 seconds then diluted 1:1 with Trypan Blue solution (Millipore Sigma), which distinguishes between live and dead cells by staining only those cells with damaged cell membranes. Cell suspensions were counted via hemocytometer to obtain live and necrotic cell counts, used to calculate percent viability.

### Determination of IC_50_ Value, GI_50_ value, and Therapeutic Index

Several metrics were employed in this study to characterize the cytotoxicity of tested compounds. To calculate the half maximal inhibitory concentration (IC_50_) value for each experiment, the absorbance value from each well was normalized against the background scan, followed by normalization to a positive and negative control using **Equation 1**, as provided in the manufacturer’s protocol.

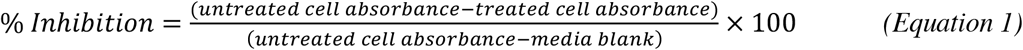

The resulting percent cell inhibition data was uploaded to GraphPad Prism9, where a nonlinear regression for Absolute IC_50_ was applied to determine the resulting IC_50_ for each compound.

For experiments testing doxorubicin, growth inhibitory GI_50_ values (i.e., concentrations for 50% of maximal inhibition of cell proliferation) were calculated for each experiment, instead of IC_50_ values. This was necessary because IC_50_ calculations could not be performed as 100% and 0% viability values were not reached within the given range of concentrations tested, a range selected based on published IC_50_ values for MDA-MB-231 cells with doxorubicin. GI50 values were calculated by plotting the percent viability against concentration for each experiment and finding the point at which the data curve intersects with 50% viability^45^.

Lastly, therapeutic index (TI) was calculated using **Equation 2** to provide a quantitative measure of the relative safety of the tested drugs,

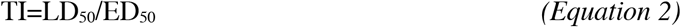

where the LD_50_ indicates the mean lethal dose of the drug on healthy cells, ED_50_ is the mean effective dose on cancer cells, and a larger TI number indicates a greater selectivity for cancer cells. For each drug, its IC_50_ value on healthy fibroblasts or erythrocytes in 2D monolayer culture was used as the LD_50_ for all calculations, while its IC_50_ value on tumor cells (for each given model cell system) served as the mean effective dose (ED_50_).

### Non-Destructive Validation of Cell Viability in Aggregates Using OCT and Trypan Blue Assay

Utilizing a recently published technique developed in our lab, we sought to validate the cell viability in tested aggregates post-SL treatment, as quantified by the CellTiter assay ^39^. Aggregates were dosed with their respective IC_50_ values as determined by serial dilutions. Concentrations selected as effective IC_50_ doses for SL-EE-D and DLSL were 125 μg/mL and 62.5 μg/mL, respectively. OCT volume scans of matured 2.5D and 3D aggregates were collected immediately prior to SL treatment, and again 12 hours post-treatment. Imaris was used as in Section 2.5 to establish volume reconstructions for each aggregate, after which the “spots” function with a 10 μm spot size (selected to match average MDA-MB-231 cell diameter) was used to identify and count the cells present within each aggregate. Three aggregates were tested for each SL and morphology combination, and three aggregates from each morphology were kept untreated as healthy controls.

To determine whether cell counts obtained via OCT/Imaris provide an estimate of the live and necrotic cells present in drugged aggregates, SL-treated aggregates were trypsinized and returned to suspension following the 12h imaging timepoint, as described in Section 2.6. Percent viability was calculated for these cell suspensions using **Equation 3**:

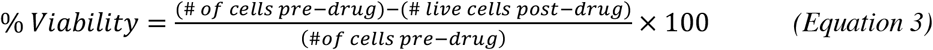

where the number of cells pre-drug for a given aggregate is obtained from OCT/Imaris, and the number of live and necrotic cells post-drug is obtained from the trypan blue assay run on that same aggregate. Comparison between these hemocytometer counts and the OCT-based cell count of *each aggregate* post-drugging could help elucidate the mechanism of cell death SLs kill cells by, i.e., any difference between these counts should indicate cells lost to apoptosis that would not appear on the trypan stain.

### Hemolytic Activity

Fresh sheep blood cells (RBCs) (Innovative Research) were used to determine the concentration of sophorolipid that causes 50% cell lysis. Red blood cells (2 mL) were washed with PBS until the supernatant became clear, using repeated centrifugation at 1000 rpm for 8 minutes to cleanse the cells. Clean RBCs (2% v/v) were mixed with increasing concentrations of either SL-EE-D or DLSL in 5% DMSO/PBS and incubated at 37°C for 1 hour. Toxicity was determined from SL-EE-D and DLSL at compound concentrations of 500, 125, 31, and 7.8 μg/mL. After incubation, RBCs were centrifuged at 1000 rpm for 8 minutes at room temperature, and 200 μL of the supernatant was transferred to a 96-well plate. Absorbance at 540 nm was recorded, with a background absorbance read at 630 nm also recorded using a BioTek uQuant Microplate Reader. Five percent DMSO/PBS + RBCs and 1% Triton X-100 (in 5% DMSO/PBS) + RBCs were used as negative and positive controls, respectively. Hemolytic activity is quantified via the HC50 value, defined as the concentration of sophorolipid that caused lysis of 50% of RBCs. Measurements were completed in triplicate. Percent inhibition was calculated using **Equation 4**^46^.

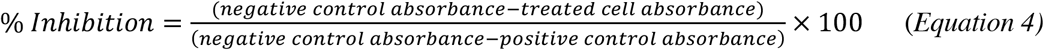

## RESULTS AND DISCUSSION

### Aggregate Morphology Characterization

The morphology of the prepared MDA-MB-231 tumor models was characterized by optical coherence tomography (OCT) imaging. Medium sized multicellular tumor spheroids (MCTSs) with diameters of 300-500 μm were targeted as they mimic several key aspects of *in vivo* tumors, such as 3D geometry, structure, and pathophysiological gradients. These features result in the formation of hypoxic cores not developed in smaller models where no gradient-related challenges are present, while avoiding the necrotic core formation seen in larger models that may no longer be representative of vascularized *in vivo* solid tumors ^47,48^. Models were produced using the liquid overlay technique with or without the addition of 2.5% Matrigel serum. Representative images of matured aggregates on Day 4 (**Fig. 2)**show significant morphologic differences between the two conditions, where Matrigel-negative samples form 2.5D disk-like aggregates while Matrigel-positive samples form truly 3D spheroids.

**Figure 2:**
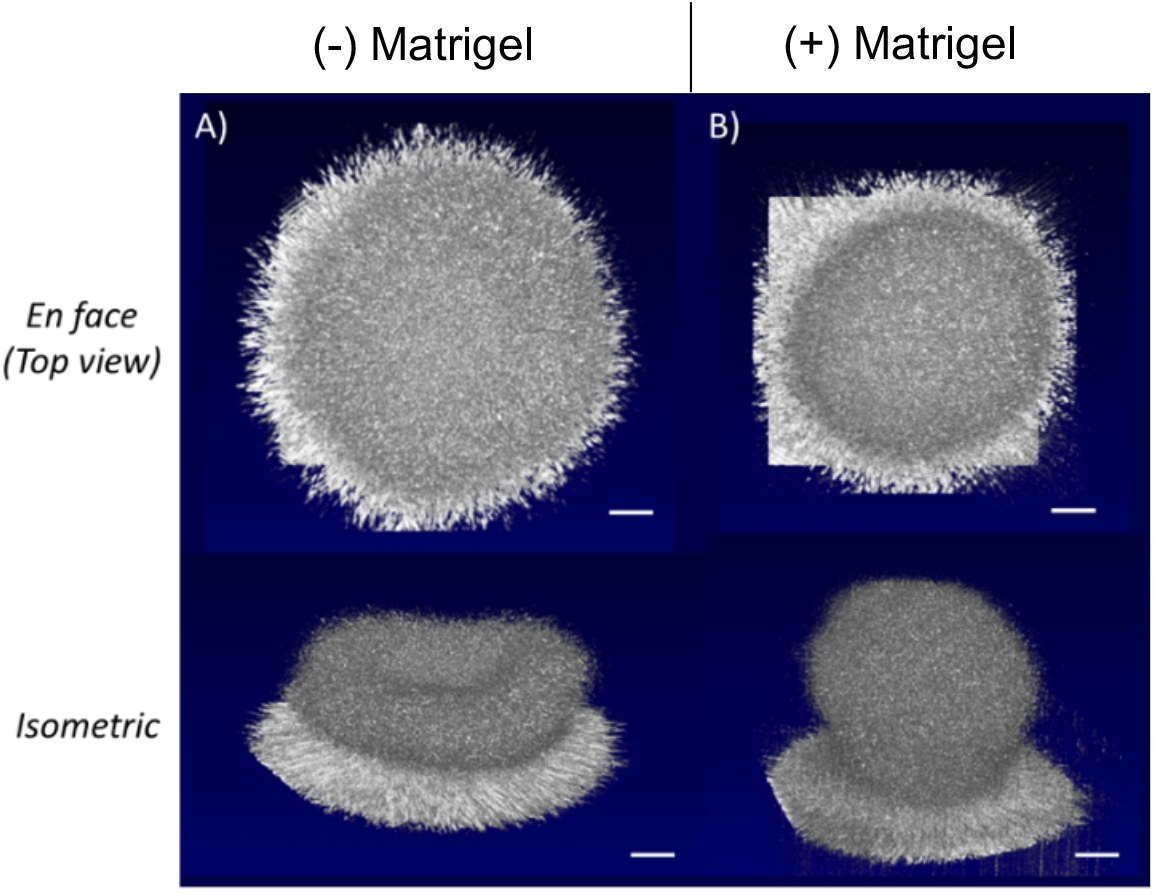
Representative Day 4 OCT images showing development of MDA-MB-231 aggregates. While both samples appear as spheroids in 2D *en face* view, (B) only those samples with Matrigel (+) formed spheroids. (A) Matrigel (-) samples formed loose, flattened, disk-like aggregates with limited compaction. Scale bars = 100 μm.

These differences in shape are quantified in **Figure 3**, where we see no significant differences in aggregate volumes, suggesting that the addition of Matrigel does not affect cell proliferation. However, we see a strikingly significant increase in sphericity with the addition of Matrigel, assessed via Student’s T-test. These geometric differences are expected to result in variations in gradient formation and, consequently, affect diffusion-associated behaviors such as cell health and drug resistance.

**Figure 3:**
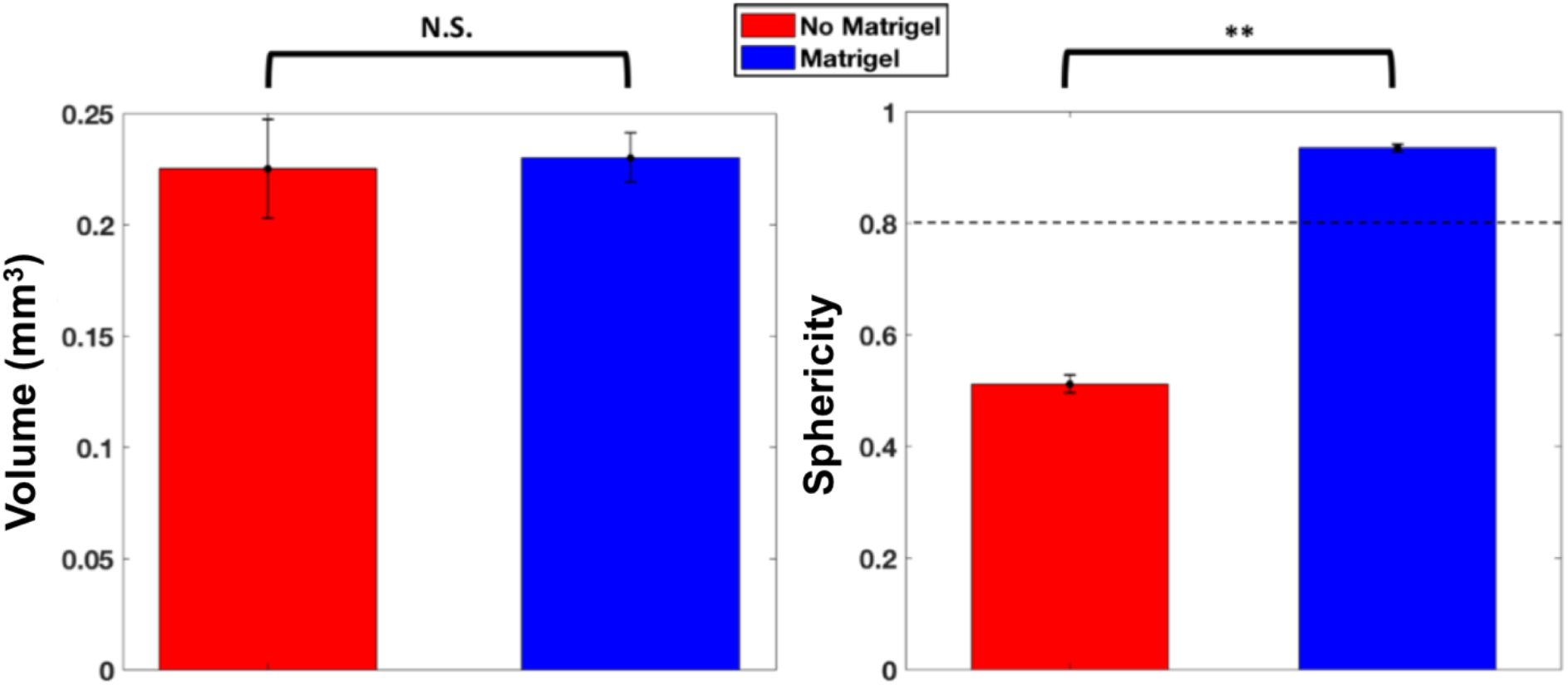
Quantification of Day 4 aggregate volume and sphericity in MDA-MB-231 cell line showed significant differences in aggregate shape despite no differences in volume. A cutoff sphericity of 0.8 was used to discern tumor spheroids from non-spherical tumor aggregates ^49^. Matrigel (+) samples produced highly spherical MCTS’s when compared with Matrigel (-) samples, which formed flatter, disk-like aggregates. (n=6) (** = p < 0.01)

### Quantification of SL Cytotoxicity

#### 2D Monolayer Results

Monolayer cultures are widely used for drug-screening due to their high-throughput capacity and overall ease. Thus, initial testing for drug toxicity was conducted on 2D cultures. Following 12 hours of drug treatment, IC_50_ values for DLSL and SL-EE-D were reported as 13.2 μg/mL and 28.9 μg/mL, respectively (**Figure 4**). These results demonstrate that, to MDA-MB-231 cells, DLSL is more toxic than SL-EE-D.

**Figure 4:**
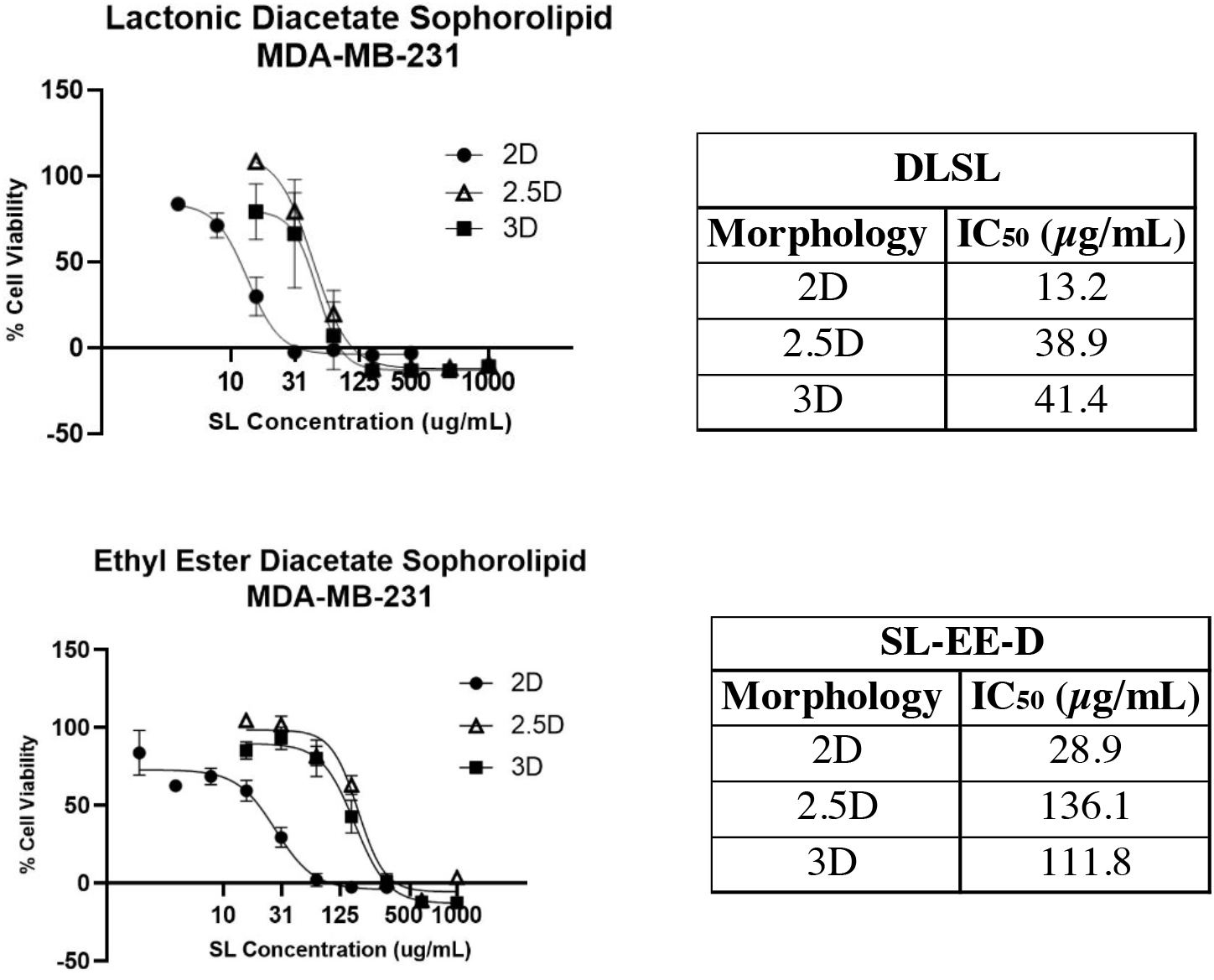
IC_50_ values obtained across conditions revealed increased drug resistance in 2.5D and 3D models compared to 2D monolayers. Both SL compounds exhibited a large decrease in cytotoxicity from 2D monolayer values to those found in 2.5D and 3D aggregate models. However, the additional of Matrigel did not cause a further reduction in toxicity on MDA-MB-231 from 2.5D to 3D aggregates, as previously hypothesized.

#### 2.5D Aggregate Results

Originally, we hypothesized that increased model dimensionality would lead to higher IC_50_ values, as this more complex morphology begins to better mimic the 3D tumor structure and gradient-behavior seen *in vivo*. To test this hypothesis, we prepared MDA-MB-231 aggregates via liquid overlay, resulting in disk-like aggregates after 4 days of maturation (**Figures 2, 3).**Following 12 hours of drug treatment on matured aggregates, we calculated an IC_50_ value of 38.9 μg/mL for DLSL (**Figure 4**). This value is approximately 3x higher than the corresponding value found in 2D culture. These results are amplified when looking at the toxicity of SL-EE-D, which showed an IC_50_ value of 136.1 μg/mL - nearly 5x higher than the dose found in 2D. In line with our hypothesis, these findings indicate that increasing aggregate/model dimensionality results in increased drug resistance. Given that these 2.5D aggregates more closely mimic *in vivo* conditions than monolayer models, these findings suggest that traditional 2D drug-screening may underestimate the required concentration of a drug, thereby underscoring the need to also evaluate drug toxicity in cell aggregate models. As an additional control experiment, the DMSO concentration present in each dose was tested in 2.5D and 3D on MDA-MB-231 cells (**Supporting Figure 3**), which showed no substantial cell death occurring due to using DMSO as solubilizing vector for SLs.

#### 3D Aggregate Results

Testing was then conducted using 3D tumor spheroids prepared via liquid overlay with Matrigel. These models have a spherical morphology and, of the models tested herein, are expected to best approximate the structure and pathophysiologic gradients seen in *in vivo* tumors. Following a 12-hour drug treatment on matured spheroids, we report IC_50_ values for DLSL (41.4 μg/mL) and SL-EE-D (111.8 μg/mL) (**Figure 4)**. These IC_50_ values are approximately 3.2x and 3.8x higher than those found in 2D, further confirming our hypothesis that increased dimensionality results in increased drug resistance. Interestingly, the increase in dimensionality from 2.5D to 3D did not result in substantially increased IC_50_ values, as was originally expected.

The IC_50_ values calculated herein for the candidate SLs demonstrate increased cytotoxicity of DLSL when compared with its modified correlate, SL-EE-D, which differs from some instances of prior literature ^6,17,18^. These findings also confirm that tumor aggregate morphology influences cellular resistance to these drugs, where increasing model dimensionality from monolayer to aggregate/spheroid was met with increased drug resistance.

### Confirmation of Cell Viability in Aggregates Using OCT and Trypan Blue Assay

Next, our recently published OCT/Imaris-based technique ^39^ was used to non-destructively verify the cell viability in 2.5D and 3D aggregates post-SL treatment at their approximate IC_50_ concentrations: 62.5 μg/mL for DLSL, and 125 μg/mL for SL-EE-D. Longitudinal Imaris cell counts, obtained from analysis of OCT images before and after a 12h SL treatment at the respective IC_50_, are shown in **Figure 5 (1A,C and 2A,C)**. Terminal trypan blue counts obtained posttreatment are shown in **Figure 5 (1B,D and 2B,D)**. Overall, both DLSL and SL-EE-D killed a significant number of cells (after 12 hours of treatment) at their IC_50_ concentrations. On average, DLSL killed 88.4 ± 3.8% of cells in 2.5D and 89.3 ± 6.3% in 3D. SL-EE-D killed 80.2 ± 9.6% of cells in 2.5D and 88.2 ± 1.2% in 3D spheroids (n=3 all cases). Imaris counts closely matched the total number of cells (live+dead) in each aggregate, as quantified by trypan blue, with an average percent difference of 11.6 ± 9.0%. Approximately one third of the cell death (29.1%) occurred via necrosis, as many dead cells were observed by hemocytometer counts. OCT confirmed that these dying cells were still structurally present. Furthermore, we hypothesize that both SLs (DLSL, SL-EE-D) also kill cells via apoptotic mechanisms. The number of total cells obtained from dissociated samples does not account for the full drop in cell viability from 0 to 12h, indicating cell death via apoptosis instead (i.e., dying cells breaking apart into pieces too small to be registered via OCT). The results of this study agree with prior literature in which both apoptotic and necrotic mechanisms of cytotoxicity have been observed when SLs were applied to cancer cells ^4,50^. Untreated 2.5D and 3D control data show no decrease in OCT-based cell count after 12h (**Figure 5: 3A,C**), indicating negligible naturally-occurring apoptosis. However, necrosis is observed in dissociated aggregates from untreated studies (**Figure 5: 3B,D**). This necrosis is likely due to aggregate morphology and the associated gradients that develop and hinder proper nutrient/waste diffusion to the center of the aggregates, thereby leading to necrosis within the aggregate core that is unrelated to drug treatment. This reasoning can help explain why more necrotic cells were observed in 3D aggregates versus 2.5D, as 3D spheroids have less favorable surface-to-volume ratios due to their increased sphericity.

**Figure 5:**
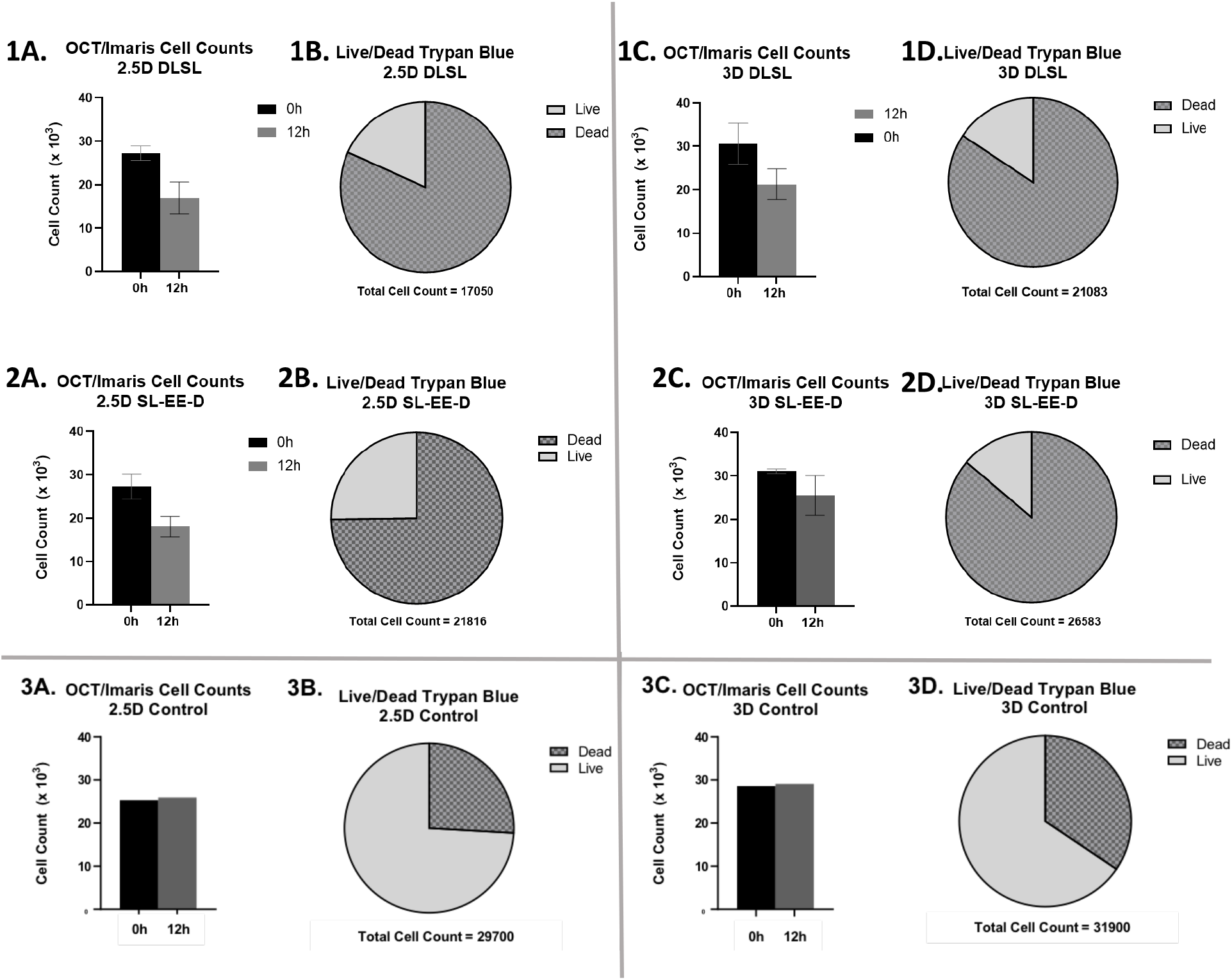
Longitudinal Imaris cell counts in MDA-MB-231 aggregates before and after 12h SL treatment at respective IC_50_ doses, presented alongside dissociated counts from the same aggregates. The number of cells present decreases for both 2.5D and 3D aggregates after 12h treatment with DLSL (1A,C) and SL-EE-D (2A,C). Dissociated cell suspensions obtained from the same aggregates post-imaging and stained with trypan blue revealed a large proportion of necrotic cells (1B,D and 2B,D). Untreated MDA-MB-231 OCT aggregate data (3A,C) shows no decrease in cell count over the 12h period, while trypan blue staining registered necrotic cells within the models (3B,D). This necrosis is likely due to a lack of media and oxygen diffusion into the center of the aggregate, and is more pronounced in 3D aggregates.

### Benchmarking Results Against a Clinically-Used Chemotherapeutic

We next sought to benchmark the performance of the candidate sophorolipids against doxorubicin, a clinically used chemotherapeutic. GI50 values were calculated for all three model types treated with doxorubicin and compared with the IC_50_ values found for the SLs. Traditional monolayer testing showed a GI_50_ value of 18.0 μg/mL (**Figure 6**), which falls within the range of IC_50_ values seen in literature for the same cell line and a similar drugging period ^40–42^. This value is slightly higher than the IC_50_ found for DLSL (13.2 μg/mL) and lower than the IC_50_ of SL-EE-D (28.9 μg/mL). In testing on 2.5D aggregates, doxorubicin yielded a GI_50_ value of 45.3 μg/mL, which is comparable to the IC_50_ found for DLSL (38.9 μg/mL) but much more toxic than SL-EE-D (IC_50_= 136.1 μg/mL). This GI_50_ value is approximately 2.5x higher than the value found in 2D, continuing the trend seen in the SLs where drug resistance increases with model dimensionality. This trend held true for inhibitory effects of doxorubicin on 3D spheroids, where the GI50 value is 37.98 μg/mL, approximately 2x higher than the value found in 2D. This value matches previously reported IC_50_ values seen for this cell line in 3D cultures ^40–42^. While doxorubicin outperformed SL-EE-D (IC_50_ = 111.8 μg/mL), our reported DLSL IC_50_ for 3D models (41.4 μg/mL) approached the doxorubicin 3D GI50 value, suggesting that DLSL may have competitive toxicity on the MDA-MB-231 cell line similar to that of the clinically used chemotherapeutic doxorubicin.

**Figure 6:**
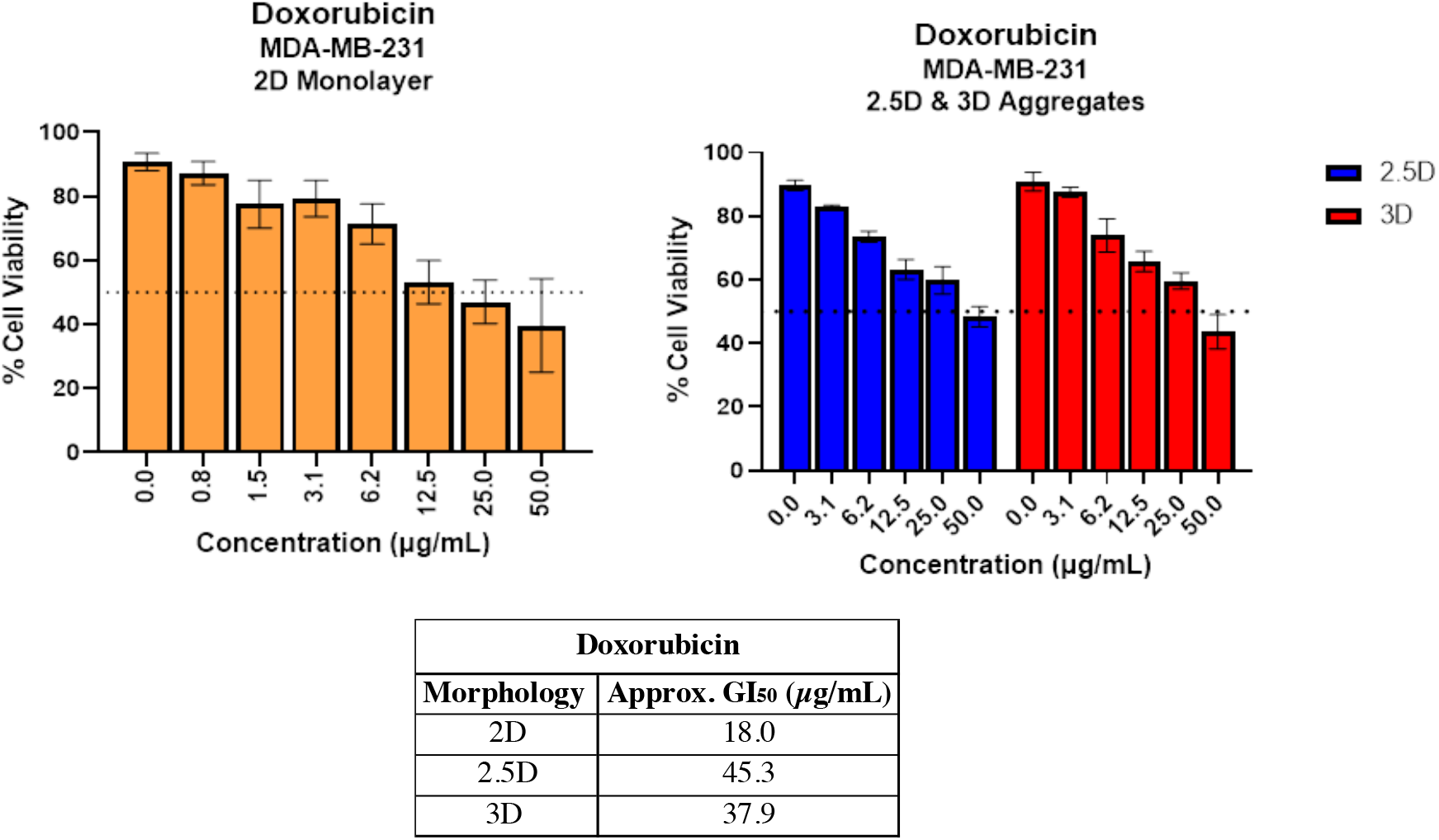
GI50 determination for MDA-MB-231 models treated with doxorubicin revealed an increase in drug resistance associated with increasing model dimensionality (from monolayer to aggregate/spheroid).

### Investigation of Cytotoxicity on Non-Cancerous Cells

#### Cytotoxicity in Fibroblasts

The cytotoxicity of the SLs on non-cancerous cells was determined to evaluate their safety. IC_50_ values were calculated for SL-EE-D and DLSL administered to fibroblasts derived from breast epithelial tissue (HDFs) (**Figure 7**), and the corresponding IC_50_ values are 32.3 μg/mL and 14.1 μg/mL, respectively. These values closely match the corresponding values obtained for the MDA-MB-231 cell line, indicating little to no selectivity of either SL-EE-D or DLSL for cancerous cells versus this specific non-cancerous cell line.

**Figure 7:**
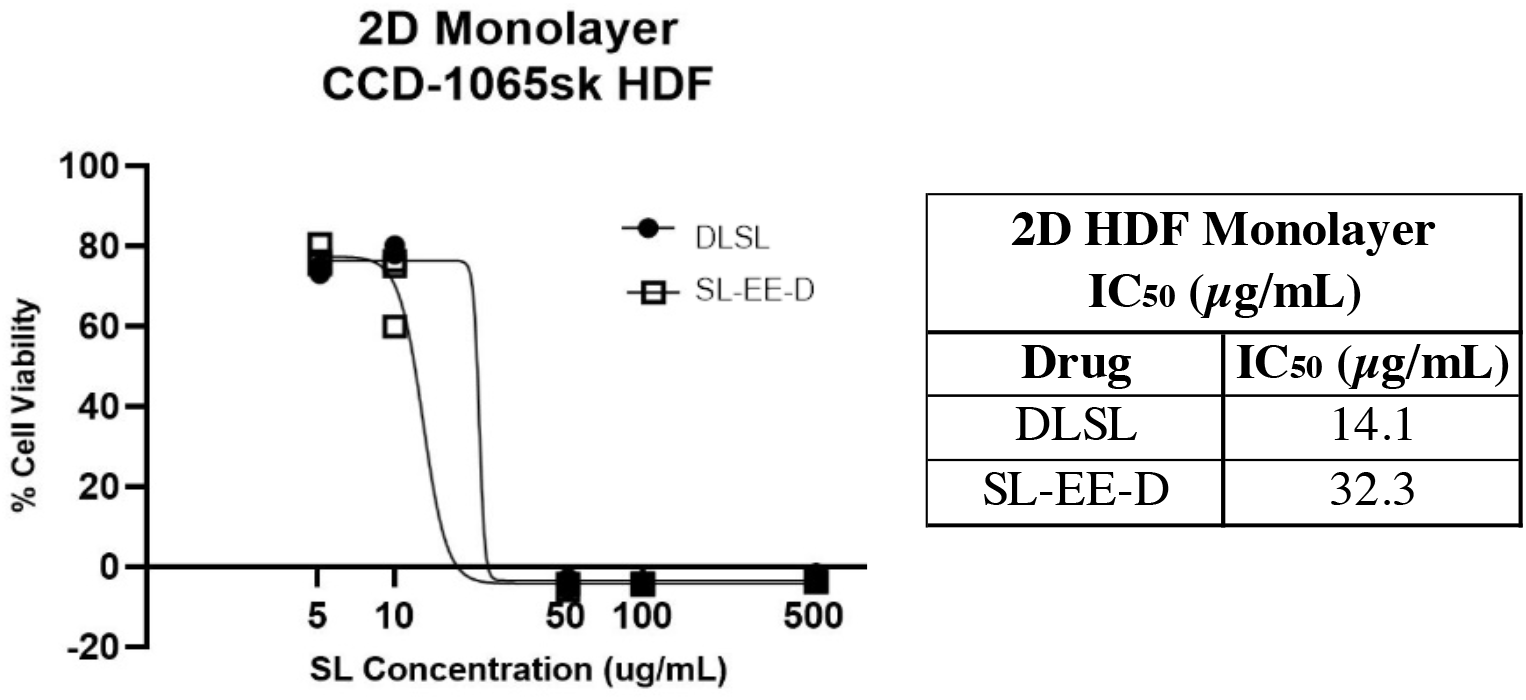
IC_50_ curves for SL-EE-D and DLSL on human dermal fibroblasts in a traditional 2D monolayer. Values closely matched those found for MDA-MB-231 cells, indicating that the SLs lack cancer selectivity and instead kill cells indiscriminately.

Next, the cytotoxicity of doxorubicin on 2D HDF monolayers was determined to establish a benchmark against which the effects of SL treatment on non-cancerous cells could be compared. We report a GI_50_ value of 29.6 μg/mL, approximately 1.6x higher than the GI_50_ dose for MDA-MB-231 cancer cells in monolayer (**Figure 8**). This relatively minor increase was expected as doxorubicin is known to possess virtually no cancer selectivity ^51^. Given that the two SL compounds studied herein showed a similar trend in IC_50_ values for cancerous and non-cancerous cells, this suggests that these SL-derived structures also possess no appreciable cancer selectivity.

**Figure 8:**
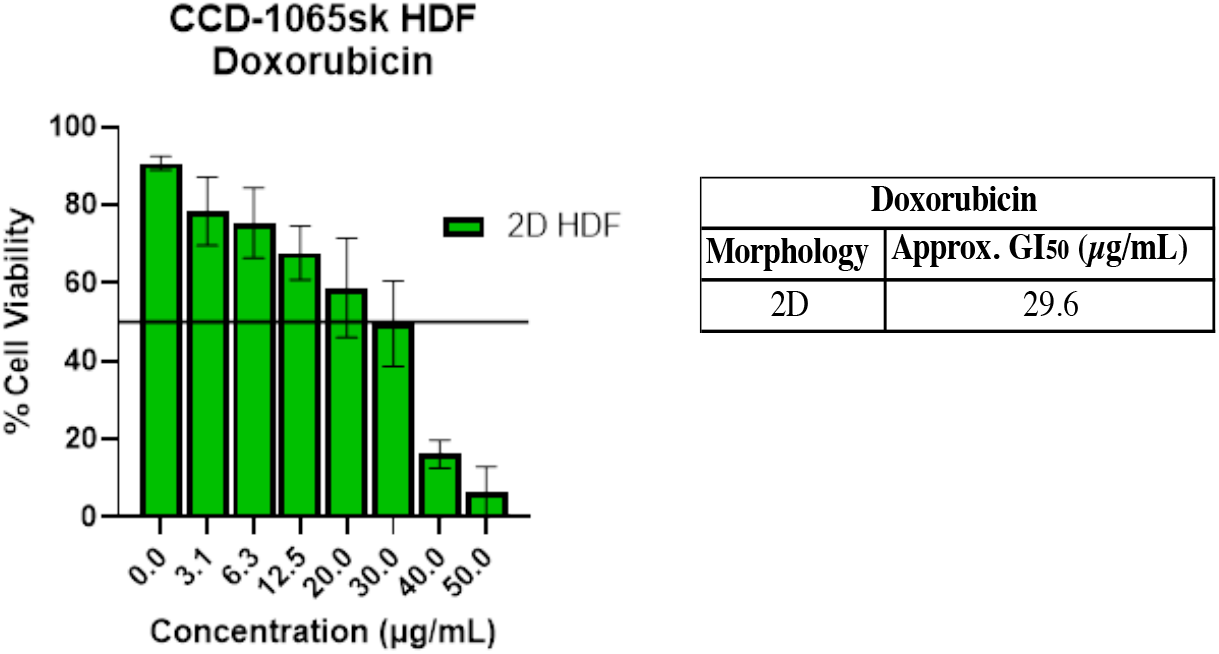
GI50 determination for doxorubicin on HDF cells in a traditional 2D monolayer. The GI50 value obtained from this study was similar to that found for doxorubicin in MDA-MB-231 cell monolayers, highlighting its general cell toxicity.

#### Cytotoxicity in Erythrocytes

Sheep red blood cells were utilized to determine erythrocyte toxicity, a common, damaging, side-effect that has previously observed for biosurfactants ^46^. It is generally concluded that a compound which causes <10% hemolysis at a given therapeutic concentration is considered nonhemolytic, while hemolyic values >25% at a given therapeutic concentration are considered to be hemolytic ^52^. Greater than 10% hemolysis occurred at SL doses of 31.3 μg/mL and above, a concentration lower that the IC_50_ doses found for thse compounds. Thus, we concluded that DLSL and SL-EE-D are hemolytic compounds. Furthermore, the HC_50_ values determined for both DLSL and SL-EE-D are each approximately 125 μg/mL (**Figure 9)**. These HC_50_ values are similar to those previously reported for other SL-analogues ^53,54^; however, this is the first report of erythrocyte toxicity for DLSL and SL-EE-D. Considering that the MDA-MB-231 IC_50_ values are lower for DLSL than for SL-EE-D at all tested morphologies (2D, 2.5D, 3D), the similarity in HC50 values suggests DLSL possesses greater cancer cell specificity than SL-EE-D. We were unable to generate an HC50 value for doxorubicin due to the autofluorescence of the compound interefering with the fluorescence of erythrocyte lysis.

**Figure 9:**
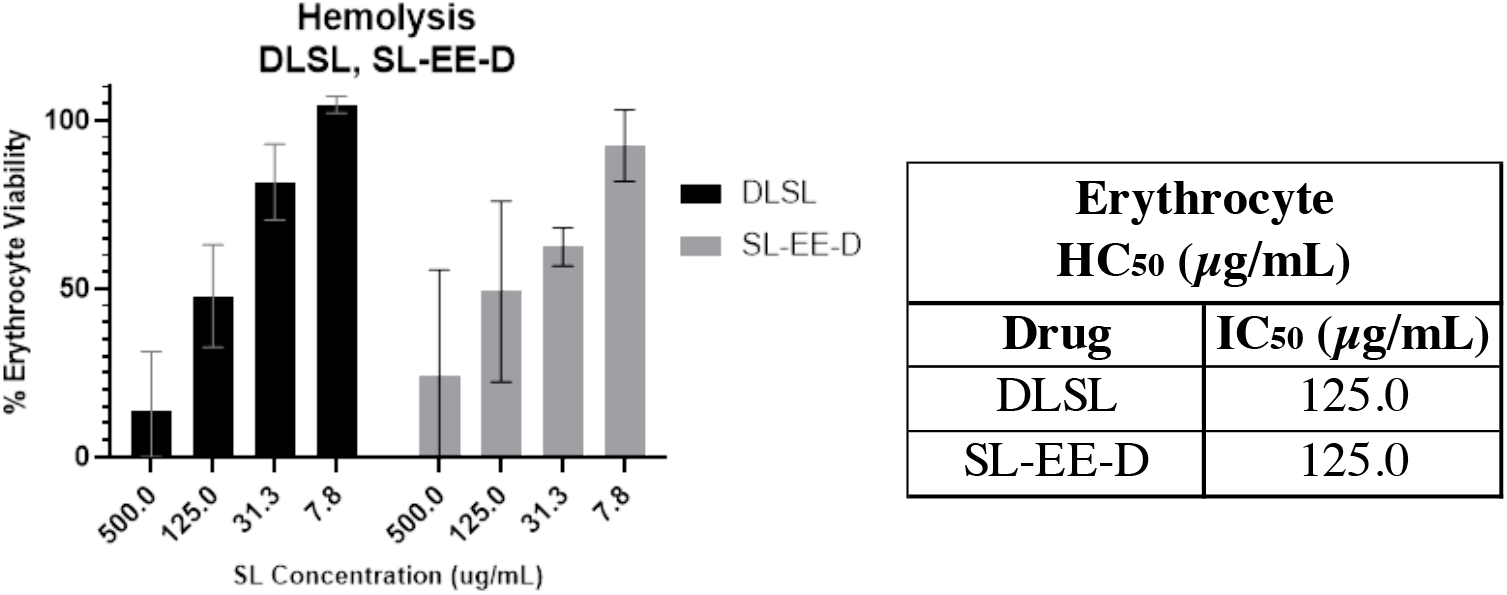
HC50 determinations for DLSL and SL-EE-D. Approximately 50% red blood cell lysis occurred at 125 μg/mL for both compounds. Data were collected in triplicate.

#### Calculation of Therapeutic Index (TI)

Lastly, therapeutic indices (TIs) for each drug at each model morphology were determined and compared. A therapeutic index provides an indication of a drug candidate’s safety by assessing the ratio of its toxicity on healthy cell lines versus the cell line of interest which, herein, is MDA-MB-231 cells. For 2D monolayers, all tested compounds had similar toxicity values towards both MDA-MB-231 and fibroblasts, resulting in low TIs (**Table 1**). TI values calculated for 2.5D and 3D MDA-MB-231 aggregates were below 1.0, likely due to the inherent difficulties in accessing these cancer cells with a drug in aggregate models. Calculation of SL TI values using erythrocytes (RBCs) rather than fibroblasts revealed improved TI values for both DLSL and SL-EE-D, indicating a selectivity for MDA-MB-231 cells over erythrocytes, particularly for DLSL.

**Table 1:**
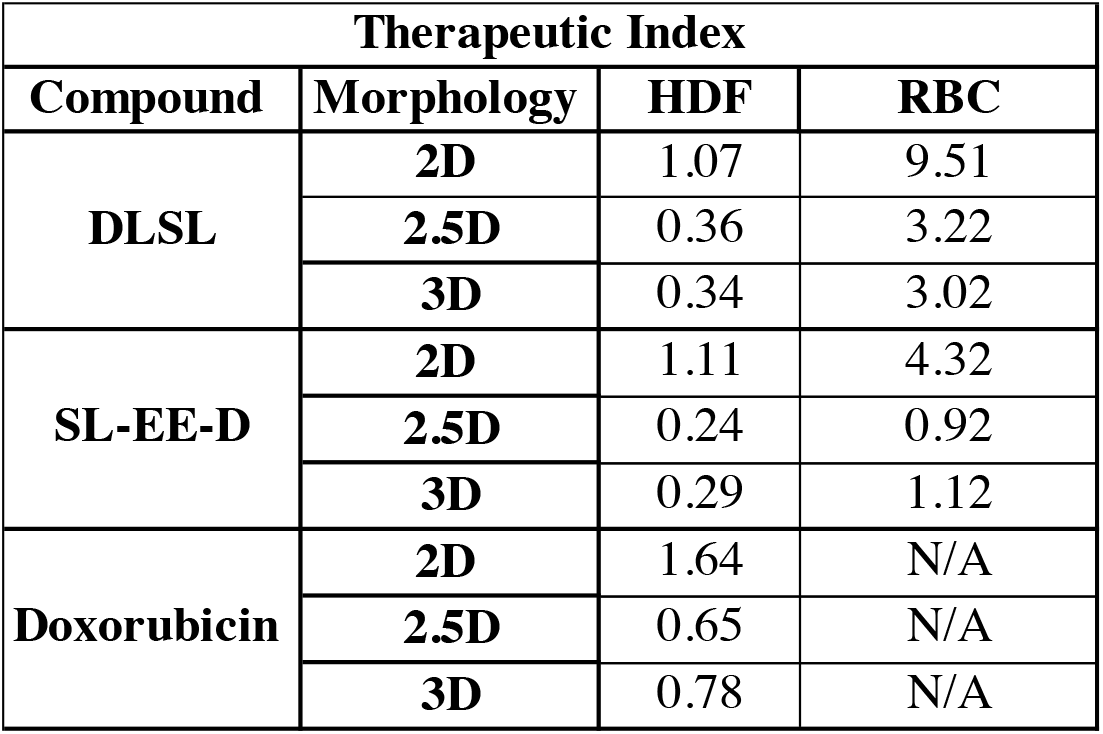
Therapeutic index as a function of model morphology for DLSL, SL-EE-D, and doxorubicin, calculated using 2D fibroblast (HDF) or erythrocyte (RBC) IC_50_ values.

## Discussion

In recent *in vitro* investigations, sophorolipids (SLs) have shown promising cytotoxic activity. Given the simplicity and lost-cost nature of their synthesis, SLs may represent an effective alternative to traditional, more costly chemotherapeutics, pending their success in preliminary drug-screening. Monolayer cell cultures are a popular choice for oncologic drug-screening due to their simplicity and high-throughput capabilities. However, a growing number of studies have shown a disconnect between these cultures and *in vivo* responses. Cells grown in monolayer do not experience gradient-related challenges, such as the diffusion of nutrients, waste, and drug, that are seen *in vivo*. Herein, we sought to expand on prior SL cytotoxic literature by introducing drug-screening results with 2.5D (disk-shaped) and 3D spheroid models, which are expected to better capture key features of the 3D tumor microenvironment. Additionally, we benchmark our findings against a clinically-used chemotherapeutic, doxorubicin, as well as testing the SL candidates on non-cancerous cells to check for cancer selectivity and safety.

Herein, we tested diacetylated lactonic sophorolipid (DLSL) and synthetically-modified sophorolipid ethyl ester diacetate (SL-EE-D) on triple negative MDA-MB-231 breast cancer cell models. Our goal was to provide preliminary data to support eventual use of these drugs as a more affordable alternative for clinical interventions. Our data show that unmodified DLSL is more lethal to these cancer cells, in all three morphologies, than SL-EE-D. The DLSL IC_50_, determined in monolayer, closely mirrored results from Riberio *et al*. with MDA-MB-231 cells (15-20 μg/mL) ^6^, and the IC_50_ of SL-EE-D is similar to that determined by Scholtz *et al*. (90% inhibition at 25 μg/mL) on Jurkat human T lymphocyte cells ^17^. Initially, testing on 2D monolayers indicated that DLSL was about twice as toxic towards the cancer cells than SL-EE-D. As we increased model dimensionality to 2.5D and 3D, DLSL became 3x as toxic towards MDA-MB-231 than SL-EE-D.

When attempting to benchmark the SLs’ IC_50_ values against those of clinically-used doxorubicin, we were challenged by doxorubicin’s autofluorescence at 450 nm. This is the same wavelength as the CCK8 cell viability dye, in which cells process tetrazolium to formazan through cell metabolism. Our attempts to use the CCK8 assay to measure the response to doxorubicin in 2D monolayers resulted in varied absorbances and high deviations between trials **(Supporting Figure 2)**. Such variations are likely due to the presence of the kit compounds, as well as increased doxorubicin autofluorescence in the wells with higher concentrations of doxorubicin present. By manually counting the live/dead cells present in doxorubicin-dosed models, we were able to compare the toxicity of this drug against that of the candidate SLs. We found that both sophorolipid compounds required similar therapeutic dose concentrations to doxorubicin in 2D monolayers, and DLSL required a similar dose to doxorubicin in both 2.5D and 3D aggregate models. These preliminary findings are promising and encourage further developmental efforts towards clinical use of these SLs.

A key finding of this study was the influence of model morphology on the drug-screening results. Upon initial model fabrication, we noted that Matrigel serum was a critical additive to achieve spherical MDA-MB-231 liquid overlay aggregate models. With an average diameter of ~500 μm, this model is expected to mimic the gradient-related behavior seen in a vascularized solid tumor. Moreover, the addition of Matrigel seemed to only affect model shape, and did not impact the aggregate volume (shown quantitatively in **Figure 3**) or cell number (data not shown). In accordance with literature, we found that 2.5D and 3D IC_50_ values were substantially higher than those found for 2D monolayers for all drugs tested, indicating higher drug resistance in these more complex models. Interestingly, while we hypothesized that 3D models would be even more resistant to drug than 2.5D models due to their more spherical shape (i.e., thicker gradients for nutrient/waste/drug transport) we found comparable IC_50_ values for these two conditions. Given that the SL molecule size is on the order of 700 Da, it is plausible that the SL molecules are too large to rely on simple diffusion through the aggregates, which would have been influenced by the different pathophysiologic gradients known to form for these models ^55^. This would be consistent with the extensive cell death observed in the trypan blue assay, indicating deep penetration of SL-EE-D and DLSL within both the 2.5 and 3D cell aggregates after only 12 hours. Overall, the substantial variation observed in cytotoxicity for a single tested drug when assayed in models of different morphologies underscores the need to include relevant cell aggregate models in drug-screening platforms. These findings point to the importance of using more physiologically-relevant three-dimensional models in preliminary *in vitro* drug-screening, given that traditional high-throughput 2D cultures may overestimate drug toxicity and lead to costly failures in clinical trials.

It is uncommon in SL literature for researchers to include testing on a non-cancerous cell lines ^4,6,17,56,57^. Herein, we included testing on fibroblasts, a cell type found in the tumor microenvironment, to assess whether our candidate sophorolipids are harmful to non-cancerous cells. We observed IC_50_ values that closely matched the corresponding values obtained for the cancer cells for both SLs. When this testing was repeated with doxorubicin, we similarly found minor differences in the GI50 values for cancerous/non-cancerous cell lines. Doxorubicin kills cells by inducing DNA double-strand breaks ^51^ and thus has notoriously low cancer cell selectivity. Given that the minor difference in GI50 values between cancerous and non-cancerous cells treated with doxorubicin matches the minor difference seen in IC_50_ values for the SLs, it appears that the two SL compounds tested also lack selectivity between the MDA-MB-231 cancer cell line and HDF’s. Testing was repeated on sheep erythrocytes, an important cell to study due to their specific plasma membrane composition and their sensitivity to oxidative stress conditions that a drug molecule can induce *in vivo*^58^. High amounts of erythrocyte hemolysis can lead to vascular irritation, phlebitis, amenia, jaundice, kernicterus, and acute renal failure ^52^. Therefore, it is important to understand how bioactive compounds interact with erythrocytes. We observed toxicity values which categorized both DLSL and SL-EE-D as cytotoxic towards erythrocytes, although we observed no appreciable difference between the HC50 value for both DLSL and SL-EE-D. Future studies in SL modification should seek to decrease SL-erythrocyte interactions, with the goal of reducing hemolytic activity.

To quantitate selectivity between cell lines, therapeutic indices (TI) for each drug were calculated for all model morphologies. The TI is a measure of drug safety as it assesses how a given drug affects neighboring, non-cancerous cells. Doxorubicin is known to have a poor TI based on numerous normal cell lines for both *in vitro* and *in vivo* environments ^59,60^. Consequently, calculated indices <2.0 using the HDF IC_50_ as the 50% effective dose (ED_50_) for all model morphologies compare well with prior studies. For the SLs, similarly low TIs were calculated using the same HDF ED_50_ value, and slightly higher TIs were calculated using the 2D erythrocyte HC50 as the ED_50_. Cytotoxicity’s and therapeutic indices from DLSL and SL-EE-D on multiple cell lines provides valuable insights into design of SL-analogues and their biological properties, particularly given that structural variants have been prepared by us ^1,16^ and others ^5,8,17,18,61^. Given the wealth of literature describing diverse chemistries that have been applied to tune SL-analogue properties^4,6,8,18,50,56,62,63^, future work with these candidate SLs should seek to explore modifications that may be applied to increase cancer cell selectivity. Additional studies should also look to confirm the results herein using an expanded set of SL-analogues, as well as testing on additional non-cancerous cells found in the tumor microenvironment (i.e., stromal cells, endothelial cells, immune cells, etc.). Additional testing with non-triple negative breast cancer cells may also reveal more specified responses within different cancer lines as opposed to the general chemotherapeutic activities of SLs explored herein. Lastly, only a few *in-vivo* studies on sophorolipid-derived drug compounds have been published ^50^. As the work on *in vitro* studies is expanded as above, *in vivo* testing with the most promising SL-drug candidates will be needed to provide better insights into pharmacokinetics, pharmacodynamics, efficacy and safety.

Lastly, we took advantage of our OCT/Imaris technique to not only non-destructively confirm our fluorescent assay results, but also to investigate the mechanism by which SLs kill cells (i.e., necrosis vs. apoptosis). The non-destructive OCT-based analyses performed herein give evidence to suggest that SLs are causing cell death by both necrotic and apoptotic pathways, a finding that has been observed previously in other SLs ^4,50^. As shown in **Figure 5**, OCT measured a decrease in cell count after 12h of SL treatment, which is hypothesized to be primarily apoptotic cell death as the cells are breaking down into structures too small to be measured by the 10 μm spots filter. Furthermore, upon dissociation of those same aggregates, we discovered most of the remaining cells undergo necrosis. These findings provide insight into the mechanisms by which SLs are killing cells, which may be used to inform their modifications towards cancer-selectivity or general increases in toxicity. Future SL studies should investigate mechanism of action through a variety of assays, including Caspase-3 and mitochondial stress to confirm apoptotic cell death, and elevated lactate dehydrogenase enzyme activity to confirm necrotic cell death ^64,65^.

Still, the techniques and methodology used herein suffer from a few shortcomings. While Matrigel is a highly effective and popular additive for 3D cell cultures, it is poorly characterized, contains many undefined growth factors, and has significant lot-to-lot compositional variability due to its mouse sarcoma origins. Thus, there is a possibility that Matrigel may be providing a yet-unknown biologic element of drug resistance to the 3D spheroid model. Future studies should seek to uncouple Matrigel’s potential biological influence from its effect on morphology. These studies may employ fabrication methods to produce 3D morphologies without Matrigel (e.g., microcapsules ^66^) or utilize Matrigel-alternatives (e.g., synthetic Matrigel ^67^, methylcellulose ^68^) to produce a 3D aggregate morphology in a more biologically mechanistic manner. Additionally, the solubility of SL drug candidates should be further modified to remove the reliance on DMSO as the vector for compound delivery. DMSO possesses high cytotoxicity at the high concentration required for initial SL solubilization, which is expected to contribute additional cell death at high SL concentrations. Furthermore, other SL-analogues may increase selectivity and hydrophilicity, without decreasing toxicity on cancerous cell lines. Lastly, given the promising results seen for the candidate DLSL tested herein, future studies should investigate additional biosurfactants for cytotoxic effects, such as modified mannosylerythitol lipids (MELs), surfactins, and rhamnolipids.

## CONCLUSION

In this work, cytotoxicity of DLSL and a novel engineered derivative, SL-EE-D, were tested on MDA-MB-231 triple negative breast cancer cells. Cytotoxicity was measured on three cell model morphologies: 2D monolayer, 2.5D aggregates, and 3D spheroids. A tandem study was completed with a clinically-used chemotherapeutic, doxorubicin, in order to benchmark SL findings. In all three drugs tested, we saw an increase in the amount of drug needed to inhibit 50% of cells from 2D models to 2.5D and 3D aggregates. The cancer-selectivity of DLSL and SL-EE-D were tested by applying the candidate drugs to both fibroblasts and erythrocytes, which revealed low therapeutic indices and minimal selectivity. To our knowledge, this is the first study to extend SL screening past 2D monolayer testing into more physiologically-relevant three-dimensional models.

## Author Contributions

The manuscript was written through contributions of all authors. All authors have given approval to the final version of the manuscript. ^These authors contributed equally.

## Acknowledgements

This study was supported by NIH R01 BRG CA207725 (DTC), NIH R01 CA233188 (DTC), and NIH NIGMS T32GM067545 (RTM). The authors would like to thank the Nuclear Magnetic Resonance Research Core in the Center for Biotechnology & Interdisciplinary Studies at Rensselaer Polytechnic Institute for their assistance with sophorolipid structure characterization. We would also like to thank Dr. Filbert Totsingan for synthesizing the sophorolipids used herein.

## SUPPORTING INFORMATION

**Table S1:**
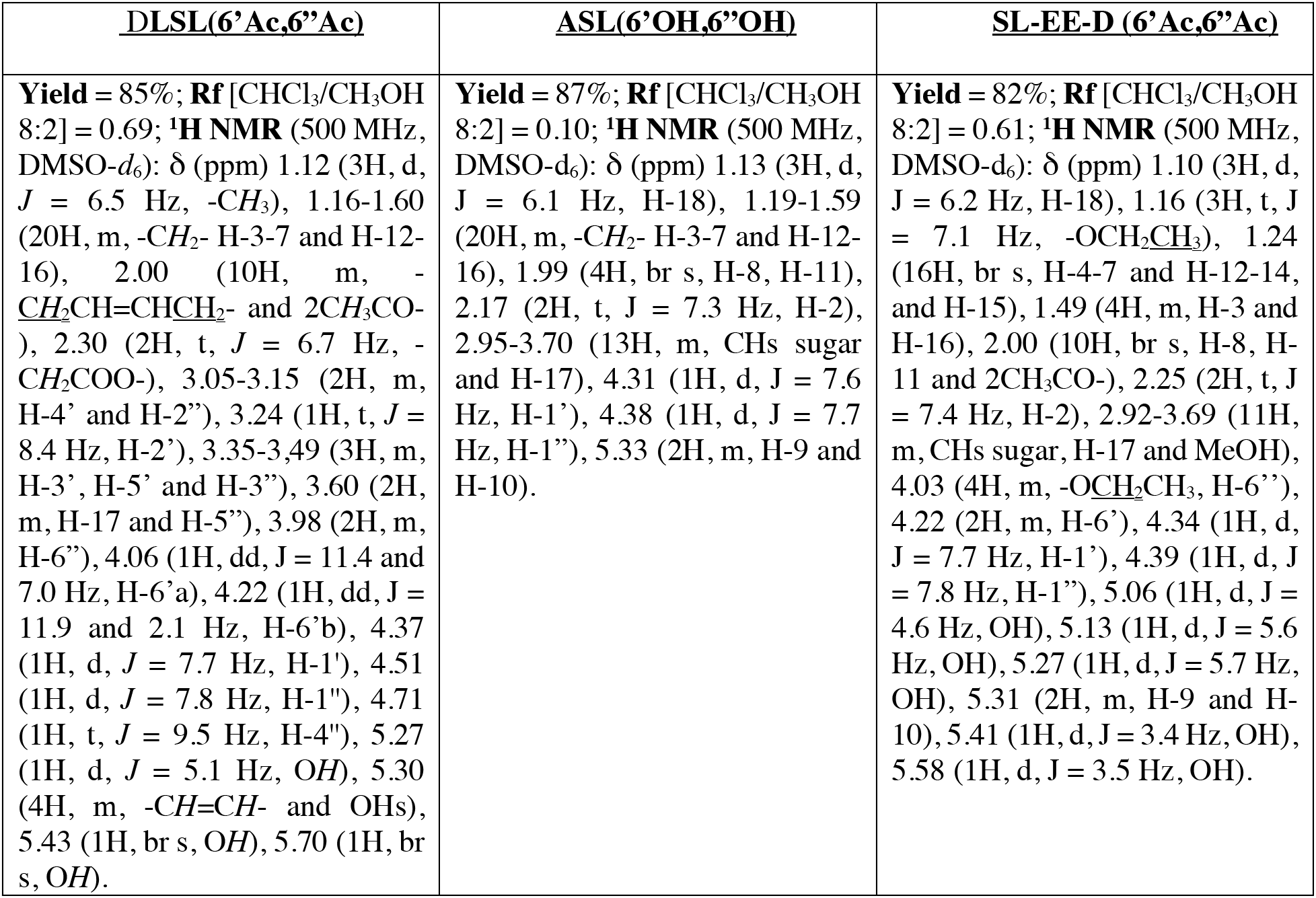
Proton NMR data from Bruker Solid State NMR

| <u>DLSL(6'Ac,6''Ac)</u> | <u>ASL(6'OH,6''OH)</u> | <u>SL-EE-D (6'Ac,6''Ac)</u> |
| --- | --- | --- |
| **Yield = 85%; Rf [CHCl<sub>3</sub>/CH<sub>3</sub>OH 8:2] = 0.69; <sup>1</sup>H NMR (500 MHz, DMSO-*d*<sub>6</sub>): δ (ppm) 1.12 (3H, d, *J* = 6.5 Hz, -CH<sub>3</sub>), 1.16-1.60 (20H, m, -CH<sub>2</sub>- H-3-7 and H-12-16), 2.00 (10H, m, -CH<sub>2</sub>CH=CHCH<sub>2</sub>- and 2CH<sub>3</sub>CO-), 2.30 (2H, t, *J* = 6.7 Hz, -CH<sub>2</sub>COO-), 3.05-3.15 (2H, m, H-4' and H-2''), 3.24 (1H, t, *J* = 8.4 Hz, H-2'), 3.35-3.49 (3H, m, H-3', H-5' and H-3''), 3.60 (2H, m, H-17 and H-5''), 3.98 (2H, m, H-6''), 4.06 (1H, dd, *J* = 11.4 and 7.0 Hz, H-6'a), 4.22 (1H, dd, *J* = 11.9 and 2.1 Hz, H-6'b), 4.37 (1H, d, *J* = 7.7 Hz, H-1'), 4.51 (1H, d, *J* = 7.8 Hz, H-1''), 4.71 (1H, t, *J* = 9.5 Hz, H-4''), 5.27 (1H, d, *J* = 5.1 Hz, OH), 5.30 (4H, m, -CH=CH- and OHs), 5.43 (1H, br s, OH), 5.70 (1H, br s, OH).** | **Yield = 87%; Rf [CHCl<sub>3</sub>/CH<sub>3</sub>OH 8:2] = 0.10; <sup>1</sup>H NMR (500 MHz, DMSO-*d*<sub>6</sub>): δ (ppm) 1.13 (3H, d, *J* = 6.1 Hz, H-18), 1.19-1.59 (20H, m, -CH<sub>2</sub>- H-3-7 and H-12-16), 1.99 (4H, br s, H-8, H-11), 2.17 (2H, t, *J* = 7.3 Hz, H-2), 2.95-3.70 (13H, m, CHs sugar and H-17), 4.31 (1H, d, *J* = 7.6 Hz, H-1'), 4.38 (1H, d, *J* = 7.7 Hz, H-1''), 5.33 (2H, m, H-9 and H-10).** | **Yield = 82%; Rf [CHCl<sub>3</sub>/CH<sub>3</sub>OH 8:2] = 0.61; <sup>1</sup>H NMR (500 MHz, DMSO-*d*<sub>6</sub>): δ (ppm) 1.10 (3H, d, *J* = 6.2 Hz, H-18), 1.16 (3H, t, *J* = 7.1 Hz, -OCH<sub>2</sub>CH<sub>3</sub>), 1.24 (16H, br s, H-4-7 and H-12-14, and H-15), 1.49 (4H, m, H-3 and H-16), 2.00 (10H, br s, H-8, H-11 and 2CH<sub>3</sub>CO-), 2.25 (2H, t, *J* = 7.4 Hz, H-2), 2.92-3.69 (11H, m, CHs sugar, H-17 and MeOH), 4.03 (4H, m, -OCH<sub>2</sub>CH<sub>3</sub>, H-6''), 4.22 (2H, m, H-6'), 4.34 (1H, d, *J* = 7.7 Hz, H-1'), 4.39 (1H, d, *J* = 7.8 Hz, H-1''), 5.06 (1H, d, *J* = 4.6 Hz, OH), 5.13 (1H, d, *J* = 5.6 Hz, OH), 5.27 (1H, d, *J* = 5.7 Hz, OH), 5.31 (2H, m, H-9 and H-10), 5.41 (1H, d, *J* = 3.4 Hz, OH), 5.58 (1H, d, *J* = 3.5 Hz, OH).** |

**Supporting mental Figure 1:**
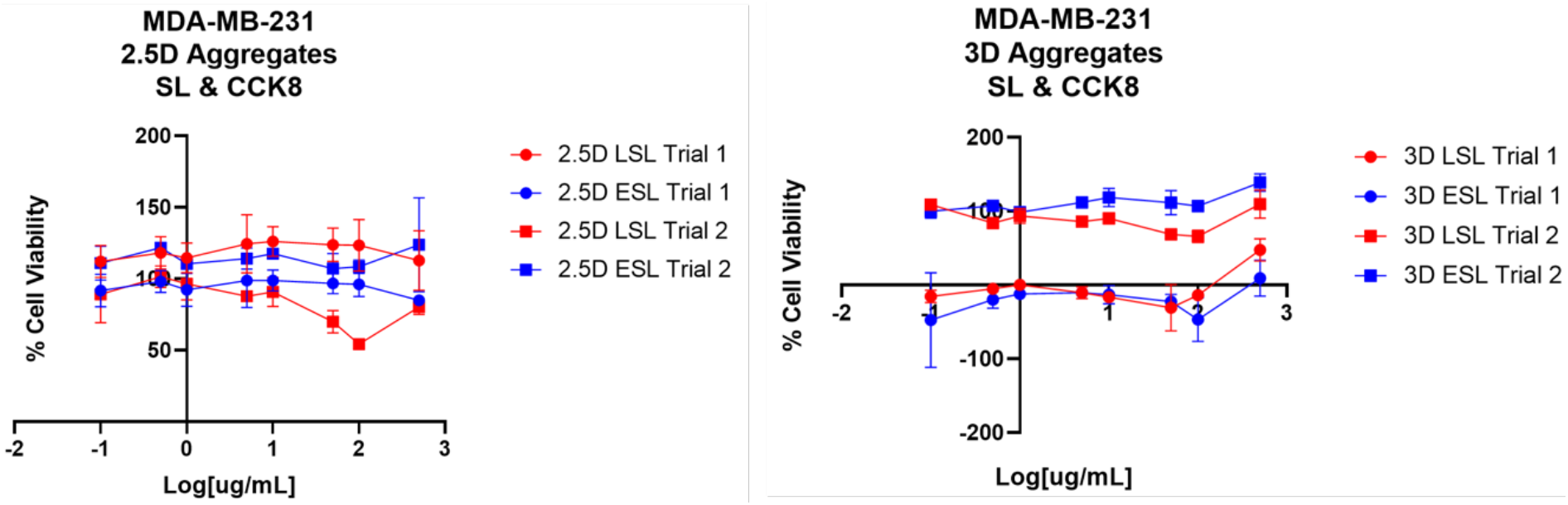
Toxicity of DLSL and SL-EE-D on 2.5D and 3D MDA-MB-231 models. Due to the random and unreproducible data values that were collected with CCK8 across multiple trials, we concluded that the assay does not provide accurate cell viability information on aggregated cell morphologies. We hypothesize that this may be due to the inability of CCK8 to diffuse into spheroids, precluding the assay from reaching inner layers of cells.

**Supporting mental Figure 2:**
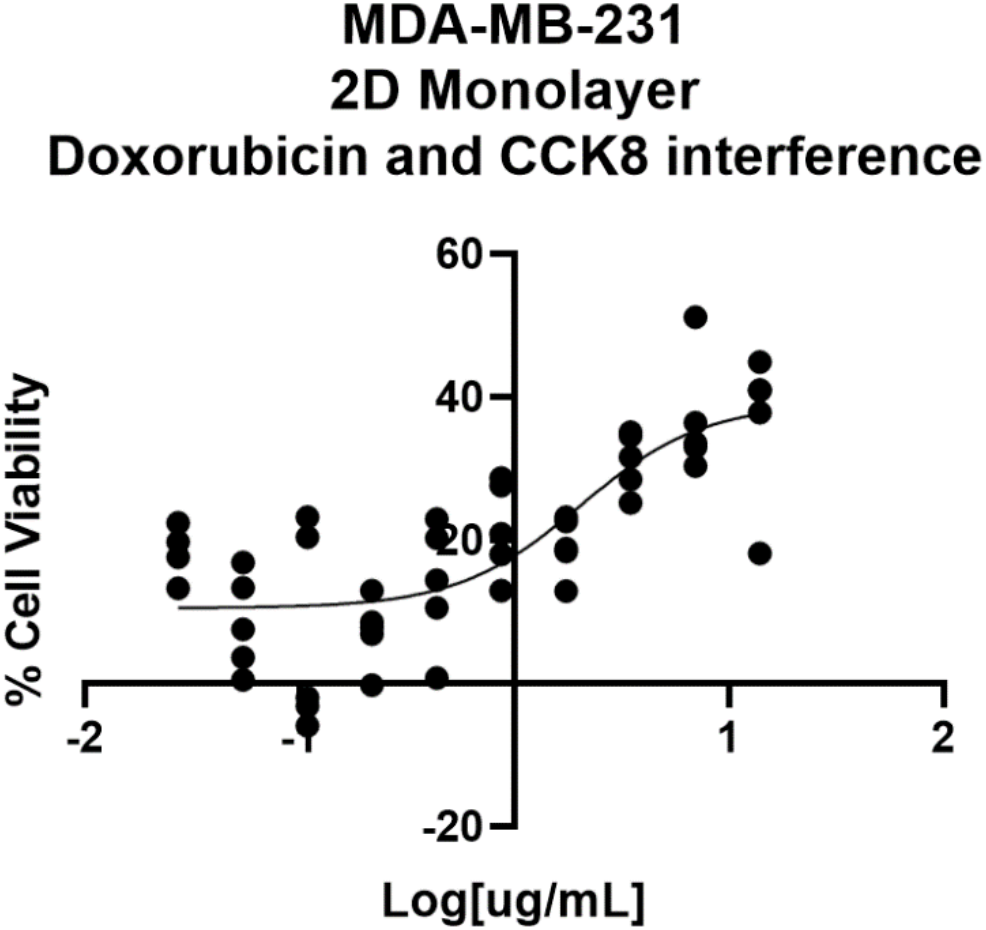
Toxicity of doxorubicin on a 2D monolayer of MDA-MB-231 cells. Doxorubicin autofluorescences at 480 nm, which is close to the 450 nm wavelength at which CCK8 is read. As a result, this assay registered increased cell viability at higher doxorubicin concentrations. Due to this interference, manual cell counting was performed to determine cell viability from exposure to doxorubicin.

**Supporting mental Figure 3:**
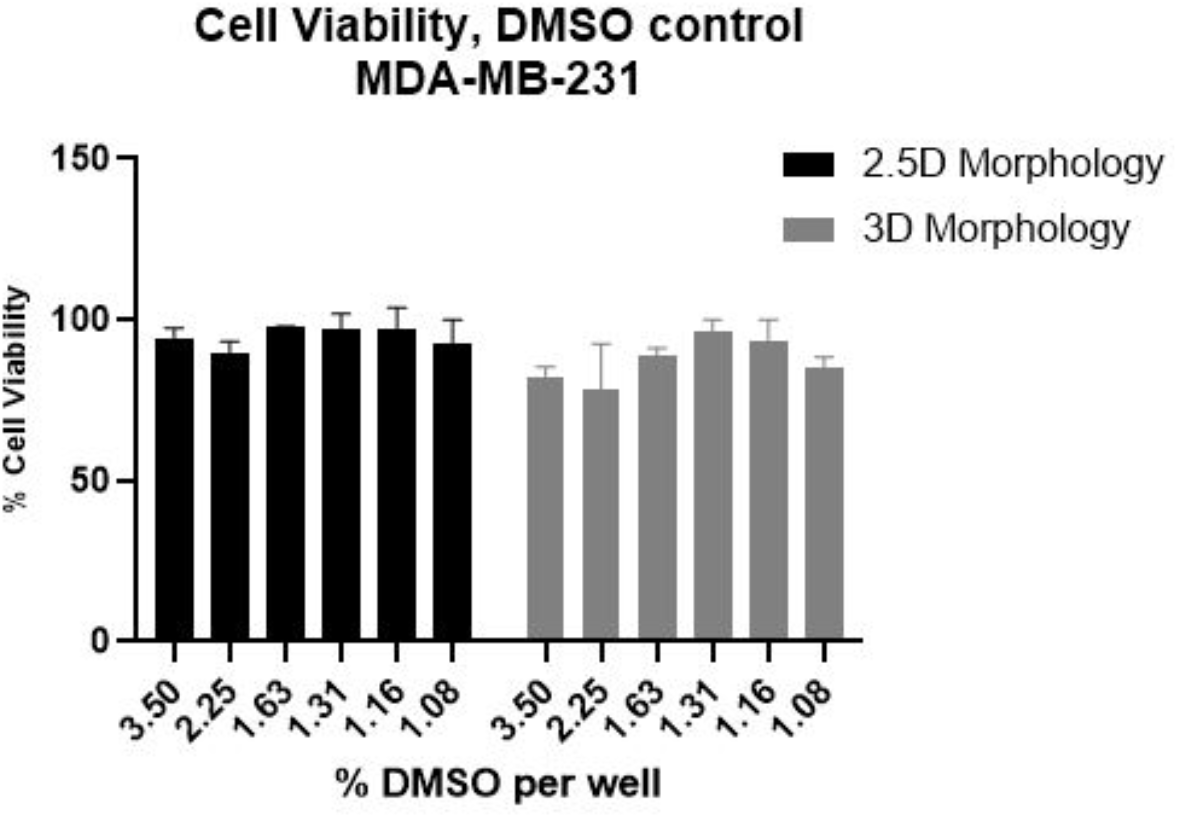
DMSO control data demonstrating no substantial cell death occurring as a result of using DMSO as solubilizing vector for SLs.

## CITATIONS

(1) Peng, Y.; Totsingan, F.; Meier, M. A. R.; Steinmann, M.; Wurm, F.; Koh, A.; Gross, R. A. Sophorolipids: Expanding Structural Diversity by Ring-Opening Cross-Metathesis. Eur. J. Lipid Sci. Technol. 2015, 117 (2), 217–228. https://doi.org/10.1002/ejlt.201400466.

(2) Singh, S. K.; Felse, A. P.; Nunez, A.; Foglia, T. A.; Gross, R. A. Regioselective Enzyme-Catalyzed Synthesis of Sophorolipid Esters, Amides, and Multifunctional Monomers. J. Org. Chem. 2003, 68 (14), 5466–5477. https://doi.org/10.1021/jo0204395.

(3) Bisht, K. S.; Gross, R. A.; Kaplan, D. L. Enzyme-Mediated Regioselective Acylations of Sophorolipids. J. Org. Chem. 1999, 64 (3), 780–789. https://doi.org/10.1021/jo981497m.

(4) Shao, L.; Song, X.; Ma, X.; Li, H.; Qu, Y. Bioactivities of Sophorolipid with Different Structures against Human Esophageal Cancer Cells. J. Surg. Res. 2012, 173 (2), 286–291. https://doi.org/10.1016/j.jss.2010.09.013.

(5) Nawale, L.; Dubey, P.; Chaudhari, B.; Sarkar, D.; Prabhune, A. Anti-Proliferative Effect of Novel Primary Cetyl Alcohol Derived Sophorolipids against Human Cervical Cancer Cells HeLa. PLoS One 2017, 12 (4), 1–14. https://doi.org/10.1371/journal.pone.0174241.

(6) Ribeiro, I. A. C.; Faustino, C. M. C.; Guerreiro, P. S.; Frade, R. F. M.; Bronze, M. R.; Castro, M. F.; Ribeiro, M. H. L. Development of Novel Sophorolipids with Improved Cytotoxic Activity toward MDA-MB-231 Breast Cancer Cells. J. Mol. Recognit. 2015, 28 (3), 155–165. https://doi.org/10.1002/jmr.2403.

(7) Miceli, T. R.; Corr, T. D.; Barroso, M. M.; Dogra, N.; Gross, A. R. Sophorolipids: Anti-Cancer Activities and Mechanisms. Bioorg. Med. Chem. 2022, 65 (February).

(8) Li, H.; Guo, W.; Ma, X. jing; Li, J. shan; Song, X. In Vitro and in Vivo Anticancer Activity of Sophorolipids to Human Cervical Cancer. Appl. Biochem. Biotechnol. 2017, 181 (4), 1372–1387. https://doi.org/10.1007/s12010-016-2290-6.

(9) Diaz-Rodriguez, P.; Chen, H.; Erndt-Marino, J. D.; Liu, F.; Totsingan, F.; Gross, R. A.; Hahn, M. S. Impact of Select Sophorolipid Derivatives on Macrophage Polarization and Viability. ACS Appl. Bio Mater. 2019, 2 (1), 601–612. https://doi.org/10.1021/acsabm.8b00799.

(10) Dierickx, S.; Castelein, M.; Remmery, J.; De Clercq, V.; Lodens, S.; Baccile, N.; De Maeseneire, S. L.; Roelants, S. L. K. W.; Soetaert, W. K. From Bumblebee to Bioeconomy: Recent Developments and Perspectives for Sophorolipid Biosynthesis. Biotechnol. Adv. 2021, 54 (June 2021), 107788. https://doi.org/10.1016/j.biotechadv.2021.107788.

(11) Gao, R.; Falkeborg, M.; Xu, X.; Guo, Z. Production of Sophorolipids with Enhanced Volumetric Productivity by Means of High Cell Density Fermentation. Appl. Microbiol. Biotechnol. 2013, 97 (3), 1103–1111. https://doi.org/10.1007/s00253-012-4399-z.

(12) Li, S.; Qian, X.; Xu, L.; Liu, S.; Xu, A.; Xin, F.; Zhou, J.; Dong, W.; Jiang, M. Biological Tailoring of Novel Sophorolipid Molecules and Their Derivatives. Biofuels, Bioprod. Biorefining 2021. https://doi.org/10.1002/bbb.2279.

(13) Borsanyiova, M.; Patil, A.; Mukherji, R.; Prabhune, A.; Bopegamage, S. Biological Activity of Sophorolipids and Their Possible Use as Antiviral Agents. Folia Microbiol. (Praha). 2016, 61 (1), 85–89. https://doi.org/10.1007/s12223-015-0413-z.

(14) Bluth, M. H.; Kandil, E.; Mueller, C. M.; Shah, V.; Lin, Y. Y.; Zhang, H.; Dresner, L.; Lempert, L.; Nowakowski, M.; Gross, R.; Schulze, R.; Zenilman, M. E. Sophorolipids Block Lethal Effects of Septic Shock in Rats in a Cecal Ligation and Puncture Model of Experimental Sepsis. Crit. Care Med. 2006, 34 (1), E188. https://doi.org/10.1097/01.CCM.0000196212.56885.50.

(15) Shah, V.; Doncel, G. F.; Seyoum, T.; Eaton, K. M.; Zalenskaya, I.; Hagver, R.; Azim, A.; Gross, R. Sophorolipids, Microbial Glycolipids with Anti-Human Immunodeficiency Virus and Sperm-Immobilizing Activities. Antimicrob. Agents Chemother. 2005, 49 (10), 4093–4100. https://doi.org/10.1128/AAC.49.10.4093-4100.2005.

(16) Totsingan, F.; Liu, F.; Gross, R. A. Structure–Activity Relationship Assessment of Sophorolipid Ester Derivatives against Model Bacteria Strains. Molecules 2021, 26 (10), 1–9. https://doi.org/10.3390/molecules26103021.

(17) C. Scholz, S. Mehta, K. Bisht, V. Guilmanov, D. Kaplan, R. Nicolosi, R. Gross. Bioactivity of Extracellular Glycolipids-Investigation of Potential Anti-Cancer Activity of Sophorolipids and Sophorolipid-Derivatives. Polym. Prepr 1998, 39, 168.

(18) Fu, S. L.; Wallner, S. R.; Bowne, W. B.; Hagler, M. D.; Zenilman, M. E.; Gross, R.; Bluth, M. H. Sophorolipids and Their Derivatives Are Lethal Against Human Pancreatic Cancer Cells. J. Surg. Res. 2008, 148 (1), 77–82. https://doi.org/10.1016/j.jss.2008.03.005.

(19) Kessel, S.; Cribbes, S.; Déry, O.; Kuksin, D.; Sincoff, E.; Qiu, J.; Chan, L. L. Y. High-Throughput 3D Tumor Spheroid Screening Method for Cancer Drug Discovery Using Celigo Image Cytometry. SLAS Technol. 2017, 22 (4), 454–465. https://doi.org/10.1177/2211068216652846.

(20) Friedrich, J.; Seidel, C.; Ebner, R.; Kunz-Schughart, L. A. Spheroid-Based Drug Screen: Considerations and Practical Approach. Nat. Protoc. 2009, 4 (3), 309–324. https://doi.org/10.1038/nprot.2008.226.

(21) Langhans, S. A. Three-Dimensional in Vitro Cell Culture Models in Drug Discovery and Drug Repositioning. Front. Pharmacol. 2018, 9 (JAN), 1–14. https://doi.org/10.3389/fphar.2018.00006.

(22) Hirschhaeuser, F.; Menne, H.; Dittfeld, C.; West, J.; Mueller-Klieser, W.; Kunz-Schughart, L. A. Multicellular Tumor Spheroids: An Underestimated Tool Is Catching up Again. J. Biotechnol. 2010, 148 (1), 3–15. https://doi.org/10.1016/J.JBIOTEC.2010.01.012.

(23) Mueller-Klieser, W. Multicellular Spheroids: A Review on Cellular Aggregates in Cancer Research. J. Cancer Res Clin Oncol. 113 (1987), 101–122.

(24) Nagelkerke, A.; Bussink, J.; Sweep, F. C. G. J.; Span, P. N. Generation of Multicellular Tumor Spheroids of Breast Cancer Cells: How to Go Three-Dimensional. Anal. Biochem. 2013, 437 (1), 17–19. https://doi.org/10.1016/j.ab.2013.02.004.

(25) Carlsson, J.; Yuhas, J. Liquid-Overlay Culture of Cellular Spheroids. Recent Results Cancer Res 1984, 95, 1–23.

(26) Rofstad, E.; Wahl, A.; Davies, C.; Brustad, T. Growth Characteristics of Human Melanoma Multicellular Spheroids In Liquid-Overlay Culture: Comparisons With the Parent Tumour Xenografts. Cell Prolif. 1986, 19 (2), 205–216.

(27) American Cancer Society. Cancer Facts & Figures 2019. Cancer Facts Fig. 2019 2019, 1–9. https://doi.org/10.1097/01.NNR.0000289503.22414.79.

(28) Nhukeaw, T.; Temboot, P.; Hansongnern, K.; Ratanaphan, A. Cellular Responses of BRCA1-Defective and Triple-Negative Breast Cancer Cells and in Vitro BRCA1 Interactions Induced by Metallo-Intercalator Ruthenium(II) Complexes Containing Chloro-Substituted Phenylazopyridine. BMC Cancer 2014, 14 (1), 1–19. https://doi.org/10.1186/1471-2407-14-73.

(29) Xiao, Z.; Morris-Natschke, S. L.; Lee, K.-H. Strategies for the Optimization of Natural Leads to Anticancer Drugs or Drug Candidates. Physiol. Behav. 2017, 176 (12), 139–148. https://doi.org/10.1016/j.physbeh.2017.03.040.

(30) Sikov, W. M.; Berry, D. A.; Perou, C. M.; Singh, B.; Cirrincione, C. T.; Tolaney, S. M.; Kuzma, C. S.; Pluard, T. J.; Somlo, G.; Port, E. R.; Golshan, M.; Bellon, J. R.; Collyar, D.; Hahn, O. M.; Carey, L. A.; Hudis, C. A.; Winer, E. P. Impact of the Addition of Carboplatin and/or Bevacizumab to Neoadjuvant Once-per-Week Paclitaxel Followed by Dose-Dense Doxorubicin and Cyclophosphamide on Pathologic Complete Response Rates in Stage II to III Triple-Negative Breast Cancer: CALGB 40603 (A. J. Clin. Oncol. 2015, 33 (1), 13–21. https://doi.org/10.1200/JCO.2014.57.0572.

(31) Nadeem, H.; Jayakrishnan, T. T.; Rajeev, R.; Johnston, F. M.; Gamblin, T. C.; Turaga, K. K. Cost Differential of Chemotherapy for Solid Tumors. J. Oncol. Pract. 2016, 12 (3), e299–e307. https://doi.org/10.1200/JOP.2015.006700.

(32) Chan, M. Cancer in Developing Countries: Facing the Challenge. 2010, p 2010.

(33) Felse, P. A.; Shah, V.; Chan, J.; Rao, K. J.; Gross, R. A. Sophorolipid Biosynthesis by Candida Bombicola from Industrial Fatty Acid Residues. Enzyme Microb. Technol. 2007, 40 (2), 316–323. https://doi.org/10.1016/j.enzmictec.2006.04.013.

(34) Van Bogaert, I. N. A.; Saerens, K.; De Muynck, C.; Develter, D.; Soetaert, W.; Vandamme, E. J. Microbial Production and Application of Sophorolipids. Appl. Microbiol. Biotechnol. 2007, 76 (1), 23–34. https://doi.org/10.1007/s00253-007-0988-7.

(35) Koh, A.; Todd, K.; Sherbourne, E.; Gross, R. A. Fundamental Characterization of the Micellar Self-Assembly of Sophorolipid Esters. Langmuir 2017, 33 (23), 5760–5768. https://doi.org/10.1021/acs.langmuir.7b00480.

(36) Ratsep, P.; Shah, V. Identification and Quantification of Sophorolipid Analogs Using Ultra-Fast Liquid Chromatography-Mass Spectrometry. J. Microbiol. Methods 2009, 78 (3), 354–356. https://doi.org/10.1016/j.mimet.2009.06.014.

(37) Koh, A.; Linhardt, R. J.; Gross, R. Effect of Sophorolipid N-Alkyl Ester Chain Length on Its Interfacial Properties at the Almond Oil-Water Interface. Langmuir 2016, 32 (22), 5562–5572. https://doi.org/10.1021/acs.langmuir.6b01008.

(38) Huang, Z.; Yu, P.; Tang, J. Characterization of Triple-Negative Breast Cancer MDA-MB-231 Cell Spheroid Model. Onco. Targets. Ther. 2020, 13, 5395–5405. https://doi.org/10.2147/OTT.S249756.

(39) Roberge, C. L.; Kingsley, D. M.; Faulkner, D. E.; Sloat, C. J.; Wang, L.; Barroso, M.; Intes, X.; Corr, D. T. Non-Destructive Tumor Aggregate Morphology and Viability Quantification at Cellular Resolution, During Development and in Response to Drug. Acta Biomater. 2020, 117, 322–334. https://doi.org/10.1016/j.actbio.2020.09.042.

(40) Pilco-Ferreto, N.; Calaf, G. M. Influence of Doxorubicin on Apoptosis and Oxidative Stress in Breast Cancer Cell Lines. Int. J. Oncol. 2016. https://doi.org/10.3892/ijo.2016.3558.

(41) Smith, L.; Watson, M. B.; O’Kane, S. L.; Drew, P. J.; Lind, M. J.; Cawkwell, L. The Analysis of Doxorubicin Resistance in Human Breast Cancer Cells Using Antibody Microarrays. Mol. Cancer Ther. 2006. https://doi.org/10.1158/1535-7163.MCT-06-0190.

(42) Tassone, P.; Tagliaferri, P.; Perricelli, A.; Blotta, S.; Quaresima, B.; Martelli, M. L.; Goel, A.; Barbieri, V.; Costanzo, F.; Boland, C. R.; Venuta, S. BRCA1 Expression Modulates Chemosensitivity of BRCA1-Defective HCC1937 Human Breast Cancer Cells. Br. J. Cancer 2003, 88 (8), 1285–1291. https://doi.org/10.1038/sj.bjc.6600859.

(43) Lutter, A. H.; Scholka, J.; Richter, H.; Anderer, U. Applying XTT, WST-1, and WST-8 to Human Chondrocytes: A Comparison of Membrane-Impermeable Tetrazolium Salts in 2D and 3D Cultures. Clin. Hemorheol. Microcirc. 2017, 67 (3–4), 327–342. https://doi.org/10.3233/CH-179213.

(44) Angello, J.; Hosick, H. Glycosaminoglycan Synthesis by Mammary Tumor Spheroids. Biochem. Biophys. Res. Commun. 1982, 107 (3), 1130–1137.

(45) Brooks, E. A.; Galarza, S.; Gencoglu, M. F.; Chase Cornelison, R.; Munson, J. M.; Peyton, S. R. Applicability of Drug Response Metrics for Cancer Studies Using Biomaterials. Philosophical Transactions of the Royal Society B: Biological Sciences. 2019. https://doi.org/10.1098/rstb.2018.0226.

(46) Kanwar, R.; Gradzielski, M.; Prevost, S.; Appavou, M. S.; Mehta, S. K. Experimental Validation of Biocompatible Nanostructured Lipid Carriers of Sophorolipid: Optimization, Characterization and in-Vitro Evaluation. Colloids Surfaces B Biointerfaces 2019, 181 (April), 845–855. https://doi.org/10.1016/j.colsurfb.2019.06.036.

(47) Jiang, Y.; Pjesivac-Grbovic, J.; Cantrell, C.; Freyer, J. P. A Multiscale Model for Avascular Tumor Growth. Biophys. J. 2005, 89 (6), 3884–3894. https://doi.org/10.1529/biophysj.105.060640.

(48) Lorenzo, C.; Frongia, C.; Jorand, R.; Fehrenbach, J.; Weiss, P.; Maandhui, A.; Gay, G.; Ducommun, B.; Lobjois, V. Live Cell Division Dynamics Monitoring in 3D Large Spheroid Tumor Models Using Light Sheet Microscopy. Cell Div. 2011, 6 (1), 22. https://doi.org/10.1186/1747-1028-6-22.

(49) Noto, A.; Raffa, S.; De Vitis, C.; Roscilli, G.; Malpicci, D.; Coluccia, P.; Di Napoli, A.; Ricci, A.; Giovagnoli, M. R.; Aurisicchio, L.; Torrisi, M. R.; Ciliberto, G.; Mancini, R. Stearoyl-CoA Desaturase-1 Is a Key Factor for Lung Cancer-Initiating Cells. Cell Death Dis. 2013, 4 (12), e947–11. https://doi.org/10.1038/cddis.2013.444.

(50) Callaghan, B.; Lydon, H.; Roelants, S. L. K. W.; Van Bogaert, I. N. A.; Marchant, R.; Banat, I. M.; Mitchell, C. A. Lactonic Sophorolipids Increase Tumor Burden in Apcmin+/-Mice. PLoS One 2016, 11 (6), 1–17. https://doi.org/10.1371/journal.pone.0156845.

(51) Pang, B.; Qiao, X.; Janssen, L.; Velds, A.; Groothuis, T.; Kerkhoven, R.; Nieuwland, M.; Ovaa, H.; Rottenberg, S.; Van Tellingen, O.; Janssen, J.; Huijgens, P.; Zwart, W.; Neefjes, J. Drug-Induced Histone Eviction from Open Chromatin Contributes to the Chemotherapeutic Effects of Doxorubicin. Nat. Commun. 2013, 4 (May). https://doi.org/10.1038/ncomms2921.

(52) Amin, K.; Dannenfelser, R.-M. In Vitro Hemolysis: Guidance for the Pharmaceutical Scientist. J. Pharm. Sci. 2006, 101 (7), 2271–2280. https://doi.org/10.1002/jps.

(53) Kumari, A.; Kumari, S.; Prasad, G. S.; Pinnaka, A. K. Production of Sophorolipid Biosurfactant by Insect Derived Novel Yeast Metschnikowia Churdharensis f.a., Sp. Nov., and Its Antifungal Activity Against Plant and Human Pathogens. Front. Microbiol. 2021, 12 (June), 1–13. https://doi.org/10.3389/fmicb.2021.678668.

(54) Tang, Y.; Jin, M.; Cui, T.; Hu, Y.; Long, X. Efficient Preparation of Sophorolipids and Functionalization with Amino Acids to Furnish Potent Preservatives. J. Agric. Food Chem. 2021, 69 (33), 9608–9615. https://doi.org/10.1021/acs.jafc.1c03439.

(55) Walzl, A.; Unger, C.; Kramer, N.; Unterleuthner, D.; Scherzer, M.; Hengstschläger, M.; Schwanzer-Pfeiffer, D.; Dolznig, H. The Resazurin Reduction Assay Can Distinguish Cytotoxic from Cytostatic Compounds in Spheroid Screening Assays. J. Biomol. Screen. 2014, 19 (7), 1047–1059. https://doi.org/10.1177/1087057114532352.

(56) Rashad, M. M.; Nooman, M. U.; Ali, M. M.; Al-Kashef, A. S.; Mahmoud, A. E. Production, Characterization and Anticancer Activity of Candida Bombicola Sophorolipids by Means of Solid State Fermentation of Sunflower Oil Cake and Soybean Oil. Grasas y Aceites. 2014. https://doi.org/10.3989/gya.098413.

(57) Jing, C.; Xin, S.; Hui, Z.; Yinbo, Q. Production, Structure Elucidation and Anticancer Properties of Sophorolipid from Wickerhamiella Domercqiae. 2006, 39, 501–506. https://doi.org/10.1016/j.enzmictec.2005.12.022.

(58) Farag, M. R.; Alagawany, M. Erythrocytes as a Biological Model for Screening of Xenobiotics Toxicity. Chem. Biol. Interact. 2018, 279 (August 2017), 73–83. https://doi.org/10.1016/j.cbi.2017.11.007.

(59) Huan, M.; Cui, H.; Teng, Z.; Zhang, B.; Wang, J.; Liu, X.; Xia, H.; Zhou, S.; Mei, Q. In Vivo Anti-Tumor Activity of a New Doxorubicin Conjugate via α-Linolenic Acid. Biosci. Biotechnol. Biochem. 2012, 76 (8), 1577–1579. https://doi.org/10.1271/bbb.120256.

(60) Zocchi, E.; Tonetti, M.; Polvani, C.; Guida, L.; Benatti, U.; De Flora, A. Encapsulation of Doxorubicin in Liver-Targeted Erythrocytes Increases the Therapeutic Index of the Drug in a Murine Metastatic Model. Proc. Natl. Acad. Sci. U. S. A. 1989, 86 (6), 2040–2044. https://doi.org/10.1073/pnas.86.6.2040.

(61) Dubey, P.; Raina, P.; Prabhune, A.; Kaul-Ghanekar, R. Cetyl Alcohol and Oleic Acid Sophorolipids Exhibit Anticancer Activity. Int. J. Pharm. Pharm. Sci. 2016, 8 (3), 399–402.

(62) Joshi-Navare, K.; Shiras, A.; Prabhune, A. Differentiation-Inducing Ability of Sophorolipids of Oleic and Linoleic Acids Using a Glioma Cell Line. Biotechnol. J. 2011, 6 (5), 509–512. https://doi.org/10.1002/biot.201000345.

(63) Akiyode, O.; George, D.; Getti, G.; Boateng, J. Systematic Comparison of the Functional Physico-Chemical Characteristics and Biocidal Activity of Microbial Derived Biosurfactants on Blood-Derived and Breast Cancer Cells. J. Colloid Interface Sci. 2016, 479, 221–233. https://doi.org/10.1016/j.jcis.2016.06.051.

(64) Strasser, A.; Vaux, D. L. Cell Death in the Origin and Treatment of Cancer. Mol. Cell 2020, 78 (6), 1045–1054. https://doi.org/10.1016/j.molcel.2020.05.014.

(65) Wang, X.; Xu, N.; Li, Q.; Chen, S.; Cheng, H.; Yang, M.; Jiang, T.; Chu, J.; Ma, X.; Yin, D. Lactonic Sophorolipid–Induced Apoptosis in Human HepG2 Cells through the Caspase-3 Pathway. Appl. Microbiol. Biotechnol. 2021, 105 (5), 2033–2042. https://doi.org/10.1007/s00253-020-11045-5.

(66) Kingsley, D. M.; Roberge, C. L.; Rudkouskaya, A.; Faulkner, D. E.; Barroso, M.; Intes, X.; Corr, D. T. Laser-Based 3D Bioprinting for Spatial and Size Control of Tumor Spheroids and Embryoid Bodies. Acta Biomater. 2019, 95, 357–370. https://doi.org/10.1016/j.actbio.2019.02.014.

(67) Aisenbrey, E. A.; Murphy, W. L. Synthetic Alternatives to Matrigel. Nat. Rev. Mater. 2020 57 2020, 5 (7), 539–551. https://doi.org/10.1038/s41578-020-0199-8.

(68) Mahboubian, A. R.; Vllasaliu, D.; Dorkoosh, F. A.; Stolnik, S. Temperature-Responsive Methylcellulose-Hyaluronic Hydrogel as a 3D Cell Culture Matrix. Biomacromolecules 2020, 21 (12), 4737–4746. https://doi.org/10.1021/ACS.BIOMAC.0C00906/ASSET/IMAGES/LARGE/BM0C00906_0008.JPEG.

